# Inability to improve performance with control shows limited access to inner states

**DOI:** 10.1101/535187

**Authors:** Marlou Nadine Perquin, Jessica Yang, Christoph Teufel, Petroc Sumner, Craig Hedge, Aline Bompas

## Abstract

Any repeatedly performed action is characterised by endogenous variability, affecting both speed and accuracy – for a large part presumably caused by fluctuations in underlying brain and body states. The current research questions were: 1) whether such states are accessible to us, and 2) whether we can act upon this information to reduce variability. For example, when playing a game of darts, there is an implicit assumption that people can wait to throw until they are in the ‘right’ perceptual-attentional state. If this is true, taking away the ability to self-pace the game should worsen performance. We first tested precisely this assumption asking participants to play darts in a self-paced and a fixed-paced condition. There was no benefit of self-pacing, showing that participants were unable to use such control to improve their performance and reduce their variability. Next, we replicated these findings in two computer-based tasks, in which participants performed a rapid action-selection and a visual detection task in one self-paced and three forced-paced conditions. Over four different empirical tests, we show that the self-paced condition did not lead to improved performance or reduced variability, nor to reduced temporal dependencies in the reaction time series. Overall, it seems that, if people have any access to their fluctuating performance-relevant inner states, this access is limited and not relevant for upcoming performance.

---

Variability is a prominent characteristic of all human behaviour. Any repeatedly performed action will show substantial variation both in how well the action is performed and in how much time is needed to perform it. This is not only true for behaviour in daily life, but can also be measured precisely during cognitive experiments. For instance, even on simple reaction time tasks featuring the same high contrast stimulus on every trial, response times (RT) show large fluctuations relative to their mean. Although variability can also have beneficial aspects (see our Discussion), it is often perceived as desirable to reduce variability as much as possible. In the lab, we seek to reduce measurement error and obtain cleaner data. In real life, we may strive to reduce variability anywhere from trivial situations, such as keeping up consistently good performance when playing darts or playing music in a band, to contexts where variability may lead to more serious consequences, such as traffic and air control.

In many situations in our everyday lives, we take it for granted that we can maximise performance by acting when we feel ready for it. For instance, in darts – and other shooting or throwing sports – players typically take a moment to concentrate and choose the ‘right’ moment to initiate an action. However, this intuition relies on two non-trivial assumptions. Let us accept that, if performance varies across time under unchanging circumstances, this has to be due to variations in some internal states. The intuition above then assumes that: 1) we can access aspects of these fluctuating internal states which are directly relevant to performance, 2) we can choose when to act accordingly in order to improve performance. The current article tests these assumptions. Specifically, we address the effects of control upon variability and performance: Participants are given a means to only start each trial when they feel ready to continue. If it is possible to have access to these performance-related internal states and to act upon this information in a useful way, the control should be an effective measure for reducing response variability and errors.

## Endogenous variability and its accessibility

In many lab-based tasks, behavioural variability can be attributed to factors inherent to the task (experimental conditions and their time order) or directly linked to the feedback (such as learning or post-error slowing, the latter referring to the phenomenon that an error on trial *n* is usually followed by a slow response on trial *n*+1; Rabbitt, 1966). However, in simple tasks, such as pressing a button in response to a visual onset, all these factors explain only a small proportion of the overall variability (Bompas, Sumner, Muthumumaraswamy, Singh & Gilchrist, 2015; Gilden 2001). The residual variance, referred to as endogenous or spontaneous, has recently received growing interest, as its properties and causes still remain largely unknown.

First, not all of this endogenous variability is random noise. Indeed, in the lab, RT on a trial is partly correlated to that on subsequent trials, and such temporal dependencies unfold on short-but also on longer-term scales (Gilden, 2001; Kelly, Heathcote, Heath & Longstaff, 2001; Wagenmakers, Farrell & Ratcliff, 2004). Similar temporal dependencies have also been found in sports performance (Gilden & Wilson, 1995; Huber, Kuznetsov & Sternad, 2016; Smith, 2003; Stins, Yaari, Wijmer, Burger & Beek, 2018; van Beers, van der Meer & Veerman, 2013). It is tempting to attribute some of this endogenous variability to familiar concepts, such as fluctuations of motivation, attention, distractibility, fatigue, arousal, or mind wandering, which may also unfold at time scales larger than one trial. It remains unclear to what extent these constructs or meta-cognitive descriptors can contribute to *explain* variability (beyond providing a label for aspects of it), but if they indeed bear some relationship to relevant internal brain and bodily states, it would be intuitive to think that these can be used to reduce variability and improve performance.

Of these meta-cognitive constructs, the concept of mind wandering in particular has received growing interest over the last decade. Mind wandering refers to the subjective report of mentally ‘tuning out’ from a task, instead focusing on thoughts that are not directly task-related (e.g. Cheyne, Solman, Carriere & Smilek, 2009; McVay & Kane, 2012). Studies designed to investigate this metacognitive construct often use the ‘probe-caught’ method (Weinstein, 2017), in which participants are interrupted during their task with a probe about their level on “on-taskness” or the amount of mind wandering they experience. Higher levels of mind wandering on these probes have been associated with higher RT variability just before the probe (Laflamme, Seli & Smilek, 2018; Seli, Cheyne & Smilek, 2013; Thomson, Seli, Besner & Smilek, 2014). This may imply that: 1) people are able to report when their thoughts are on- or off-task, 2) this subjective report bears some relation to their recent performance, and as such, that 3) participants can access some aspects of their internal fluctuating states (but see Discussion). However, even if relevant information were available on these internal states, the extent to which people could use it to reduce their own variability or improve their upcoming performance is rarely addressed.

Mind wandering is tightly linked to the more traditional cognitive concept of attention, although the exact relation remains unclear. A possible distinction may be the level of awareness: While mind wandering requires some form of awareness (even if this awareness is ‘post-hoc’), as it is primarily a subjective mental state, this may not necessarily be the case for episodes of low task-focus (also known as lapses of attention). Like mind wandering, attention has been linked to behavioural variability. It has been said that “attention quenches variability” (Masquelier, 2013, p.8), as more attention and higher predictability correlate with lower variability on both a neuronal and a behavioural level (Cohen & Maunsell, 2009; Ledberg, Montagnini, Coppola & Bressler, 2012; Mitchell, Sundberg & Reynolds, 2007). Lapses of attention are typically suspected when RTs are very slow, but also when they are extremely short (so called *‘anticipations’*, Cheyne et al., 2009), the combination of which leads to increased variance. Yet another link between attention and mind wandering is that patients with Attention-Deficit and/or Hyperactivity Disorder (ADHD) are typically thought to suffer from lapses of attention, and have been reported to show higher variability as well as higher spontaneous mind wandering in comparison to a non-clinical population (Seli, Smallwood, Cheyne & Smilek, 2015; Shaw & Giambra, 1993; see Kofler et al., 2013 for a meta-analysis; see Tamm et al., 2012 for a review).

Although in the literature, there is a strong reliance on attention and mind wandering as causal factors for behavioural variability, it remains unclear what these concepts exactly refer to, how they relate to each other, and how they are exactly linked with variability. Still, it seems intuitive that variability in performance is caused by fluctuations in some underlying brain and body states. Our main question here is whether such states are accessible to us and whether we can act upon this information.

## Reducing variability with control?

The potential use of control in reducing variability may seem intuitive when looking at sports. For instance, when thinking about playing darts, there is the implicit assumption that people have access to some internal states as well as means to act upon them – leading them to throw the darts when they ‘feel ready for it’. When playing darts, people may feel that they have the ability to wait until they feel fully attentive to the board and to throw on this exact moment. Within this framework, taking away one’s ability to self-pace their darts game should deteriorate performance. However, while the origins of variability in dart throwing have been of interest in sports and movement psychology (e.g., Smeets, Frens & Brenner, 2002; Stins et al., 2018; van Beers et al., 2013), this specific prediction seems not to have been empirically tested so far. For now, it remains unknown what constitutes this feeling of ‘being ready’, how it links to our internal states, and whether it actually influences performance.

Unlike in a game of darts, in a traditional experimental psychology paradigm, timing of actions is carefully planned and controlled: The time from each trial to the next (‘inter-trial interval’; ITI) is determined externally, either by an absolute timing or by a jitter with a fixed range and mean. Being in an unfavourable internal state when the trial starts or when the target appears could lead to poor performance on that trial. Thus, giving participants control over the timing of the task – by letting them start a new trial whenever they feel ready for it, thus creating a ‘self-paced’ task – may enable them to reduce their variability, by preventing extreme RT and errors.

To our knowledge, this is the first study that compares a self-paced condition (in which participants determine their own ITI) to ‘forced-paced conditions’ (in which the ITIs are calculated from the self-paced ITIs) as a means to reduce variability and improve performance. Kelly et al. (2001) investigated the effects of ‘self-pacing’ versus ‘forced-pacing’ on temporal structure of choice RT. However, in their study, the ‘pacing’ refers to the maximum response time allowed after stimulus onset – as a means to manipulate the difficulty levels of the conditions. While participants were thus given some form of control (they could allow themselves more or less time to respond, and this triggered the onset of the next trial), their design does not address the question of the current research – whether control to *start* a trial can help improve on-going performance and reduce variability. Specifically, Kelly et al. (2001) investigated differences in temporal structure of choice RT series on a four choice serial RT task, and found that RT series in the ‘self-paced’ condition (in which response time was unlimited) indeed showed less long-term dependency (i.e. being closer to white noise) compared to the ‘forced-paced’ conditions (a ‘fast’ version, in which the maximum response time was the mean of the self-paced condition, and a ‘slow’ version, in which the maximum response time was the mean plus two standard deviations of the self-paced condition). They also looked at performance (but not variability), and found that mean RTs were higher in the self-paced condition. However, because both of their forced-paced experiments consisted of a fixed ITI, while the self-paced condition was not fixed but rather differed from trial to trial, findings may therefore be attributed to differences in the variability of the ITIs.

Our aim is to test whether participants can access their fluctuating performance-related internal states and have the means and will to act upon these to improve their performance (referred to as *Hypothesis 1* or *H1* throughout the article). The alternative hypothesis (*Hypothesis 2* or *H2*) is thus that people either have no access to performance-related internal state, or no will to act accordingly or no means to improve their performance as a result. In most of the tests below, but not all, *H2* is equivalent to the null hypothesis. Because of this, we use Bayesian statistics throughout the article in order to assess the evidence in favour of *H2* even when it is equivalent to a null finding.

In our first experiment, we test *H1* within its intuitive framework: With a darts-based task. Highly motivated participants played a game of darts both with and without control over when they could throw. If *H1* is true, participants should be able to use the control in the darts game to obtain higher and less variable scores compared to when they have no control. Under the alternative hypothesis (*H2*), no decrement in performance would be expected when control is taken away from participants.

The second experiment uses a computer-based design consisting of two different tasks (easy and hard visual detection tasks) – in order to converge two different literature fields (fast action selection and visual perception). In these two tasks, participants are given control or no control over the ITI. The goal of the second experiment is three-fold. First, to replicate and to generalise our findings from Experiment 1 over various forced-paced control conditions. Second, to test another two predictions of H1 related to the RT and ITIs (which were not available in Experiment 1), namely that 1) long ITIs should be associated with better performance, and 2) RT series in the self-paced condition should show fewer temporal dependencies. Third, Experiment 2 allows for closer examination of the self-paced ITIs themselves, to see how participants might use the control they are given.

## Experiment 1 – Testing the use of control in a darts task

### Rationale and Predictions

The first experiment involved participants throwing darts in self-paced and in forced-paced manners. There are multiple advantages to using darts: 1) there is a clear intuitive link between darts, control, and insight into perceptual-attentional states, as discussed in the Introduction, 2) similar to laboratory experiments, darts involves performing the same action over and over again, 3) unlike laboratory experiments, people typically can play darts for a good deal of time without getting bored, 4) the darts board can be set up with a scoring system that allows for a measure of performance and, 5) participants can easily understand what constitutes ‘good’ and ‘bad’ performance (an explicit monetary reward was used to reinforce this) and 6) participants would be motivated to get the best performance, and thus, motivated to take advantage of the control when offered to (motivation was also independently assessed via a questionnaire).

The darts task consisted of two conditions: 1) the Self-paced condition, in which participants throw the dart whenever they feel ready, and 2) the Forced-paced condition, in which participants are instructed to throw in a forced-paced (but comfortable) manner according to a tone. To further increase motivation, social competition in pairs (Tauer & Harackiewicz, 2004) and a random lottery reward system (Cubitt, Starmer & Sugden, 1998) were used – both of which have been shown to be effective for increasing motivation in participants.

If participants can use the control in the Self-paced condition to throw at the ‘right’ moment (H1), this should result in higher average scores (darts closer to bull’s eye) and lower variability compared to the Forced-paced condition. However, if they cannot use the control (H2), performance and variability should be similar under both conditions.

It is important to note that the measure of variability does not stand on its own. All in all, we are looking for *consistently good* performance – meaning that the variability should be interpreted in light of the performance and not as a sole measure of performance (particularly since reducing variability was not part of the instructions – participants were instructed to perform well, but were not explicitely told to be consistent). For example, a lower mean score in combination with lower variability would indicate that participants are consistently worse, not better. Instead, consistently good performance would be reflected in the combination of higher scores and lower variability.

Because a self-paced darts task may be more familiar to participants compared to throwing darts to a tone, we analysed the scores over block, to examine if potential practice effects would be different between the conditions even after the initial training phase. An additional analysis was conducted on the scores of the last block only, as these blocks should be the least affected by practice effects.

## Methods

### Participants

In total, 22 participants (14 female, 21-39 years, *M*_*age*_ = 25.3 years) with normal or corrected-to-normal vision were tested. All participants were right-handed. They were paid £8 as a base rate for participation (excluding reward). At the end of the experiment, all the participants filled the Intrinsic Motivation Scale (IMI; McAuley, Wraith & Duncan, 1991). One participant was excluded from analyses because of a low motivation score (less than half of the possible highest score of 144, making her a statistical outlier) – leaving 21 participants for analyses, whose average reported IMI score was 111 (*SD* = 8.37, ranging between 91-124). Bayesian statistics were used and, rather than carrying on recruiting until our Bayes Factor reached a pre-determined cut-off, we judged the null result sufficiently convincing after the first wave of recruitment (moderate evidence against H1, Figure 1C) and turned to a better-controlled experiment allowing richer data to be collected (see Interim Discussion 1). The study was approved by the local ethics committee.

**Figure 1.**
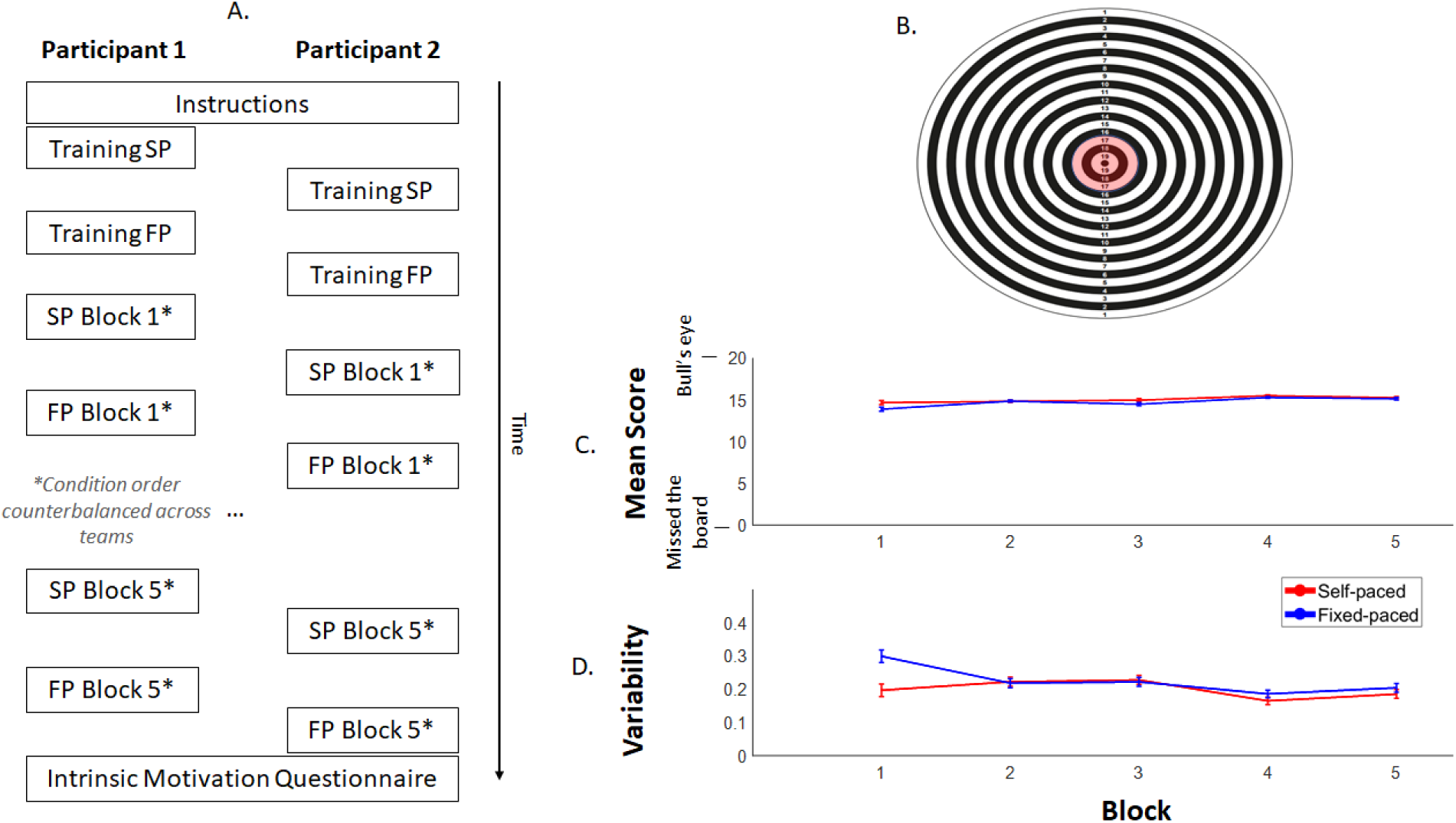
***A***. Structure of Experiment 1. Participants played darts in pairs. In turns, they would first perform a training of the Self-paced (SP) condition, to get used to throwing the darts, and then a training of the Forced-paced (FP) condition, to learn the rhythm of the tones. Next, they would play five blocks of each condition, with the order of the conditions being counterbalanced across pairs. Each block consisted of twelve trials. At the end, participants filled in the Intrinsic Motivation Scale. ***B.*** Target sheet (A2-sized) covering the dartboard, indicating the points for each ring, from the outer ring (1 point) to the bull’s eye (20 points). Trials in which participant scored within the four most inner circles (in red) qualified for reward. ***C.*** Average score per dart on each block of twelve trials on the Self-paced (red) and Forced-paced (blue) condition. ***D.*** Average coefficient of variation (CV = SD_score_ / Mean_score_) on each block. Error bars show the within-subject standard error. There was no effect of condition (arguing against Hypothesis 1).

### Materials

The darts game was played using a 45.1cm by 45.1cm Winmau dartboard and twelve nylon shafted Winmau darts. The board was covered with printed target sheets with 20 black and white rings (see Figure 1B). The scores of the rings went up by one point per ring with the most outer ring being worth one point and the bull’s eye (inner circle) being worth 20 points. For each participant, four target sheets were collected: one for each training condition, and one for each experimental condition.

The experiment was run using MATLAB 9 (The MathWorks, Inc., Release 2016a) and Psychtoolbox-3 (Brainard, 1997; Kleiner, Brainard & Pelli, 2007; Pelli, 1997). Tones were presented over Logitech s-150 USB digital speakers (Logitech, Lausanne, Switzerland). During the experiment, participants’ scores were recorded using a scoring sheet.

### Design

Both tasks had two conditions: Self-Paced and Forced-paced. In the Self-Paced condition, participants were instructed to throw the darts one by one in their own tempo – providing them with control over the timing of the action. In the Fixed condition, participants were instructed to throw in a fixed rhythm. On each trial, they heard three tones:

1) A low tone to indicate ‘ready’, on which participants were instructed to pick up a dart – followed by 1000ms of silence.
2) A low tone to indicated ‘steady’ – on which participants were instructed to get into a throwing position – followed by 1500ms of silence.
3) A low tone to indicate ‘go’ – on which participants were instructed to throw the dart – followed by 1000ms of silence before the next trial started.

The timing of the Forced-paced condition was based on pilot data, designed to ensure that the Forced-paced condition would be comfortable for participants and would have similar block durations as in the Self-paced blocks. In the main experiment, the Self-paced blocks turned out to have lower block durations on average than the Forced-paced blocks – and as such, any potential poor performance in the Forced-paced condition could not be due to the participants not having enough time to throw.

### Procedure

Participants were tested in pairs, with the full session lasting about an hour. Figure 1A shows the complete timeline of a session. First, participants were given instructions on the structure and rules of the experiment. Then, they chose the order in which they played. In total, each participant completed two training blocks (one for Self-paced and one for Forced-paced) and ten experimental blocks (five for Self-paced and five for Forced-paced). Each block consisted of twelve trials. Participants would throw six darts, have a short break in which the experimenter would get the darts off the board, and then throw six more darts.

Participants alternated their game between each block: The first participant would play one block of one condition, next, the second participant would play the same block of the same condition, then the first player would play one block of the other condition, and finally the second participant would play one block of the other condition. Between each block, the experimenter switched the paper targets on the board. After both participants had finished a block, the total scores would be compared, and one participant was named the winner of that block. For the experimental blocks, half of the pairs started with the Self-paced condition and half of the pairs started with the Fixed-paced condition to counterbalance for an order effect. Training blocks were exempt from counterbalancing: To get used to throwing with the darts, all pairs started with the Self-paced training, followed by the Fixed-paced training.

The dartboard was hung at a height of 153 cm. Participants stood at 152cm from the board. A line of masking tape was put on the floor to indicate where they had to stand exactly. The six darts were laid out in a row on a table next to them. At the beginning of each run of six darts, the experimenter told the participant when to start and pressed a key on the keyboard to record the start time. At the end of the run, the experimenter again pressed a key, to obtain the total time of the run.

At the end of the game, the experimenter drew a random trial number and checked for both participants if they were eligible for the extra reward: If a participant had a score of seventeen or higher on that trial (four most inner circles), he/she would receive £5 extra, but if the score was sixteen or lower, he/she would only receive the base rate of £8. This cut-off was chosen to get a 20% chance of winning the reward (based on pilot data).

## Results

Training trials were excluded from all analyses. Average scores and CV (coefficient of variation, equal to standard deviation of score divided by the mean score) were calculated over the five blocks of twelve trials. All statistics were Bayesian and conducted in JASP (JASP Team, 2017), using equal prior probabilities for each model, and 10000 iterations for the Monte Carlo simulation.

First, to assess the overall effect of condition, Bayesian 2×5 Repeated Measures (RM) ANOVAs were performed on the scores and CV, with condition and block as factors. Figure 1C and D show the means and CV of the Self-and Forced-paced conditions over the five blocks. Using a Bayesian RM ANOVA with condition and block as factor, each possible model was compared to a null-model (Table 1). On both mean score and CV, data were more likely under the null-model than under the model with condition (BFM0 = .75 for means and .82 for CV – meaning that the data are respectively (1 / .75) 1.33 times and (1 / .82) 1.22 times more likely under the null-model than under H1), but evidence was in the indeterminate range. Overall, for both measures, the model with only block as a factor performed better than any model that included the condition factor and/or its interaction with block. When looking at the BF_inclusion_ of each factor – which reflects the average of all models that include that factor – block is above 1. All in all, these results suggest that there is only an effect of practice.

**Table 1.**
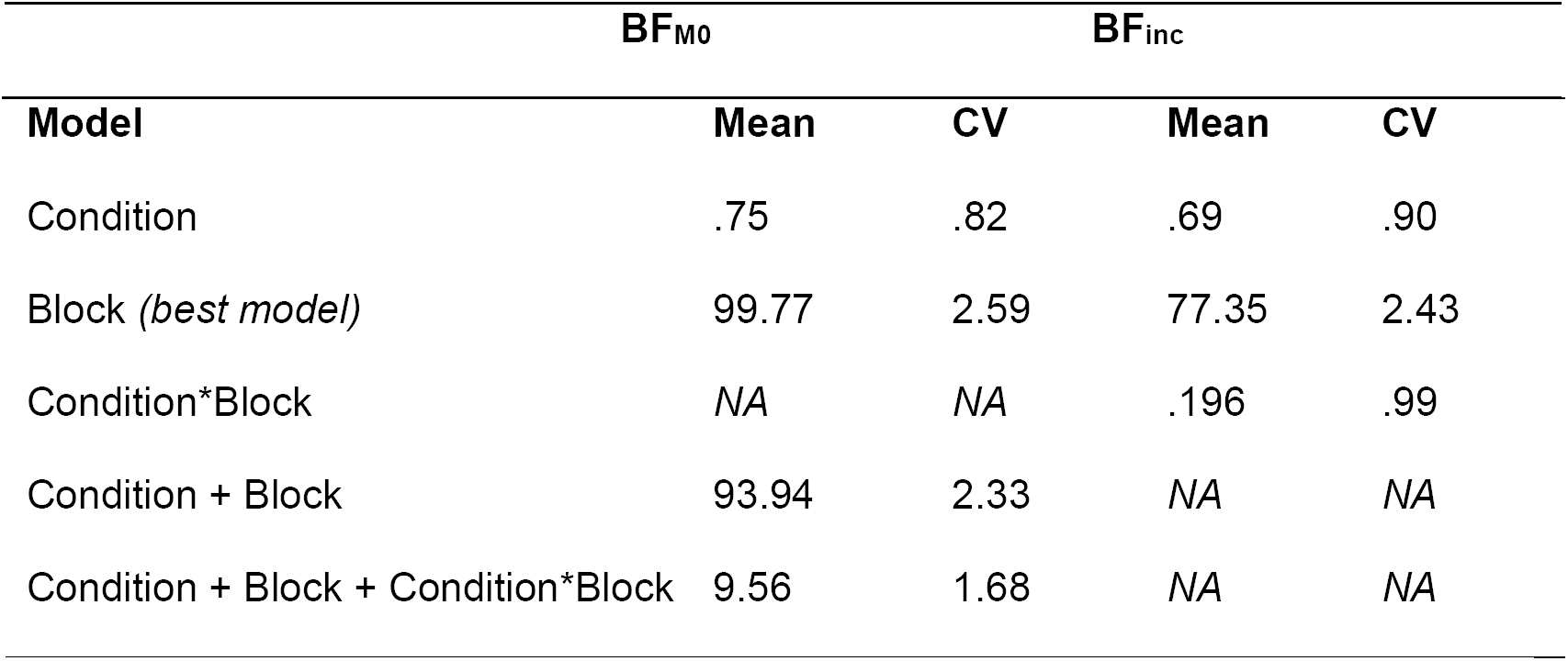
Statistical outcomes of the Bayesian RM ANOVAs on the mean score and the coefficient of variation (CV) of score, using condition and block as independent factors. BF_M0_ reflect the Bayes’ Factors of the comparison between each model to the null-model.

| **Model** | **Mean** | **CV** | **Mean** | **CV** |
| --- | --- | --- | --- | --- |
| Condition | .75 | .82 | .69 | .90 |
| Block (*best model*) | 99.77 | 2.59 | 77.35 | 2.43 |
| Condition*Block | NA | NA | .196 | .99 |
| Condition + Block | 93.94 | 2.33 | NA | NA |
| Condition + Block + Condition*Block | 9.56 | 1.68 | NA | NA |

Secondly, to directly assess H1 versus H2, a Bayesian Paired t-test was conducted on the scores and CV of the last block of both conditions. Because H1 specifically predicts an improvement in the self-paced condition, while H2 could predict either no difference or worsened performance, the t-test was conducted one-sided. The Bayesian paired one-sided t-tests on the last block showed moderate evidence for H2 over H1 on average score (BF_21_ = 3.05) and similar conclusion on variability (BF_21_ = 2.33) – again suggesting there was no improvement with control.

Lastly, to test for a potential effect of condition order, two-sided Bayesian Independent Samples t-tests were conducted on the scores and CV on each block for each condition, using order as grouping variable. There was no evidence for such confound on either measure (BFs ranging between .39-.84).

### Interim discussion 1

Overall, we found no evidence for a benefit of control in Experiment 1; throwing darts in a self-paced manner did not lead to higher performance or reduced variability compared to throwing on a fixed rhythm. When looking only at the scores of the last block, in which participants are most familiar with both of the conditions, we found moderate evidence against an effect of control – suggesting that if there was any initial benefit of throwing in a self-paced manner, it was due to unfamiliarity with the paced protocol.

Note that the interpretation of the intra-individual variability may not be straightforward, as it only reflects the variability of the raw scores, and not the variability of the position on the board. For example, imagine that a participant throws one dart on ring 14, one on 15, and one on 16. These darts could be close together or scattered over the board, but the measured variability would be the same in either case. However, the variability together with the mean performance reflects what is most important: Whether or not control can lead to consistently good performance.

One limitation of Experiment 1 is that the two conditions are quite different from each other in terms of timing – similar to the design of the Kelly et al. (2001) study. Rather than using a standard fixed pace for each participant, it would be more appropriate to use participants’ own self-paced timings as a control condition. However, this is difficult to achieve in the darts experiment, as we did not possess a way to easily measure RTs. Therefore, we aimed to replicate our findings in a computer-based experiment, to have more flexibility over the timing of the forced-paced condition. This experiment will also allow us to measure temporal dependencies in RT series. Another limitation of Experiment 1 is the relatively low number of trials per participant (60), while the complex manual action is sensitive to trial-to-trial motor noise – the combination of which may lead to decreased statistical power. Due to its traditional set-up, Experiment 2 has a larger amount of trials and is therefore more sensitive to capturing the mean score and intrinsic variance.

Because the self-paced ITIs are recorded in Experiment 2, it allows us to examine these in more details, to see what potential strategies participants may use while handling the control. The analyses for Experiment 2 are therefore split into two parts, with the first part being focused on the effects of control, and the second on characteristics of the ITI.

## Experiment 2 – Testing the use of control in two computer-based tasks

### Rationale

The second experiment involved two different tasks: An easy, action-oriented task (a rapid action selection task) and a difficult, perception-oriented task. The action-oriented task is easy to perform, and therefore participants will immediately notice their own errors. The perception-oriented task involves near-threshold stimuli tailored to produce 25% errors on average. These two tasks aim to cover two different literatures: The mind wandering literature, in which it is common to use simple tasks that are highly familiar and repetitive in nature (see for example: Cheyne et al., 2009; Seli et al., 2013; Thomson et al., 2014) – as these types of simple tasks are well-suited for inducing mind wandering (Cheyne, Carriere & Smilek, 2006; Giambra, 1995) – and the literature on perception and noisy decision making (see for example: Ergenoglu et al., 2004; De Graaf et al., 2015; Romei, Gross & Thut, 2010; Romei, Rihs, Brodbeck & Thut, 2008), in which it is common to use visually-challenging detection tasks.

In both tasks, a target appeared either on the left or the right side of the screen on each trial, and participants were asked to indicate on which side the target appeared. Both versions of the task consisted of four conditions: 1) the Self-paced condition, in which participants manually start each trial themselves, 2) the Fixed condition, in which the ITI is the same for each trial, 3) the Replay condition, in which the ITIs of the self-paced condition are Replayed in the exact same order, and 4) the Shuffled replay condition, in which the ITIs are Replayed in a shuffled order. These four conditions were inspired from Marom & Wallach (2011), although their research question was different from ours. Because the self-paced ITIs differ from traditional ITIs on multiple aspects, the three forced-paced conditions were chosen such that each of them allows for comparison with the self-paced condition over a different aspect (see Table 2 for an overview), and allows us to determine the influence of control on performance and variability, as well as on dependencies in the data over time. The Fixed condition is most similar to the forced-paced condition of Experiment 1 as well as to traditional experimental designs, and allows for predictability of trial-onset. The Replay condition is an exact replica of the self-paced ITIs and thus has the same variability, but it may therefore contain temporal dependencies. In the Shuffled Replay condition, these time structures are removed. This means that to ascribe any result to an effect of control in the Self-paced condition, the result has to be consistent over all three of the forced-paced conditions.

**Table 2.** Summary of the four different conditions and the main characteristics of the ITIs. The three forced-paced conditions (shaded in grey) allow for comparison to the Self-paced condition on these different characteristics.

| Aspect | Condition |  |  |  |
| --- | --- | --- | --- | --- |
|  | Self-paced | Fixed | Replay | Shuffled replay |
| Control over ITI | Yes | No | No | No |
| Predictability of trial onset | Yes | Yes | No | No |
| Variability of ITI | Yes | No | Yes | Yes |
| Time structure in ITI | Yes* | NA | Yes* | No |

\* See Section “Characterising the self-paced ITIs in the computer-based tasks” and Supplementary Table 1.

Table 3 gives an overview of the two different hypotheses as well as the corresponding empirical predictions and findings over Experiment 1 and 2. To contrast our hypotheses, we investigate the effects of task-control and spell out four empirical tests, the predicted outcomes of which differ across hypotheses. We compare the performance (RT and accuracy), intra-individual variability (CV of RT), and serial dependencies in the self-paced condition with the three forced-paced conditions. With Test 1 and Test 2, we aim to replicate the findings from Experiment 1 – that participants cannot use the control to improve their performance and reduce their variability. Test 3 and 4 offer additional tests of *H1*, examining the impact of long ITIs on performance and contrasting temporal dependencies in RT series across conditions.

**Table 3.** Summary of the two alternative hypotheses and their respective predictions over the two experiments. Green shading indicates those predictions that were supported by the data in the present article. Evidence favoured H2 (people have no access to performance-relevant inner states or no will/means to act upon it) over Hypothesis 1 (people have access to performance-relevant inner states and will plus means to act upon it).

| Empirical predictions Experiment 1 | Hypothesis 1 | Hypothesis 2 |
| --- | --- | --- |
| Improved performance and reduced variability in Self-paced | Yes | No |
| Empirical predictions Experiment 2 |  |  |
| 1. Improved performance and reduced variability in Self-paced | Yes | No |
| 2. Reduced extreme RTs | Yes | No |
| 3. Performance following long self-paced ITIs is: | Better | Worse |
| 4. Reduced temporal dependencies in Self-paced | Yes | No |

Unlike Experiment 1, Experiment 2 comes with the possibility of recording self-paced ITIs, and therefore using these in designing the forced-paced conditions. In order to record and replay the self-paced ITIs, the Self-paced condition will have to come first. Although we showed in Experiment 1 (which allowed for counterbalancing of conditions) that order did not matter, a concern might be that participants could continue to show training effects in the Self-paced condition – which could mask differences between the conditions. To anticipate, we found no evidence for such training effects, making it unlikely that the results are explained by condition order.

#### Test 1. The effect of control on performance and variability

First, we aimed to replicate the results from Experiment 1, i.e. that having control does not lead to improved performance or reduced variability. For both tasks, we calculated for each condition: 1) mean reaction time, 2) percentages of errors, and 3) coefficient of variation of RT (CVRT). Again, if we do not have access to our own internal performance-related rhythms or have no means to act upon them, having control over the timing should *not* lead to lower RT means, lower error rate, or lower CVRTs.

Just as in Experiment 1, it is important to not interpret the measures individually. When investigating RT and accuracy, good (or poor) performance is not just indicated by each of them separately, but by the combination of the two. For example, a reduced mean RT with an increased error rate is not indicative of improved performance as such, as it could also reflect an adjustment of speed-accuracy trade-off. Here, we use the EZ-diffusion model to investigate this in more detail (see Test 1b). Furthermore, we are again looking for *consistently good* performance – meaning that the variability can only be interpreted in combination with performance.

##### Test 1b. The effect of control on performance as EZ-diffusion model parameters

The EZ-diffusion model was used to disentangle strategy adjustments from true performance improvements. The EZ-diffusion model is based on the drift-diffusion model (DDM; Ratcliff, 1978), which is a computational model for two-alternative forced choice tasks – in which participants have to make a choice between two options (in this case, ‘left’ or ‘right’). The model assumes that evidence accumulates between two boundaries, each representing one response option, until one of them is reached, which initiates the corresponding response.

The EZ-diffusion model is a simplified version of the DDM (Wagenmakers, Van der Maas & Grasman, 2007), which uses calculations rather than a fitting procedure. It provides three parameters: 1) drift rate (v), which reflects the rate with which evidence is gathered (or in other words, how quickly information is processed), 2) boundary separation (α), which reflects a response criterion (or in other words, reflects how much evidence is needed before an action can be initiated), and 3) non-decision time (T_er_), which reflects the time spent on any processes but decision making (such as sensory and motor execution). Improved performance may be reflected in higher drift rates and/or in lower non-decision times, while differences in speed-accuracy trade-offs may be reflected in the boundary separation.

#### Test 2. Reduced extreme RTs

Extreme RTs are considered the main hallmark of attentional lapses. Even if participants are not able to reduce their mean RT, they may still be able to use the control to avoid these extreme reaction times. To test for this, the number of very long *and* very short reaction times (likely anticipations) was calculated for each condition and each participant. Under the intuitive framework of *H1*, in which people can wait for the ‘right’ moment to perform, participants in the Self-paced condition should be able to delay the start of the next trial until their attentional lapse has passed, leading to a reduced amount of extreme reaction times. But under *H2*, there should be no difference between the conditions.

#### Test 3. The effect of longer self-paced ITIs

To get more insight into potential ways participants may have used the ITIs, we tested whether longer ITIs reflected moments when participants waited for a more optimal moment to initiate the trial. ITIs were devided ‘regular’ and ‘long’, and the mean reaction time, coefficient of variance, and accuracy were calculated on the ‘regular ITI’-trials and on the ‘long ITI’-trials. If participants can effectively make use of the control – i.e. if they can use these longer breaks to wait until they feel ready to continue (*H1*) – their performance should increase and their variability decrease on the trials with long self-paced ITIs compared to trials with regular self-paced ITIs.

Alternatively, participants may simply show fluctuating good and poor modes of responding throughout the experiment over which they have no control, similarly affecting both RT and ITIs. If this were the case, these long ITIs may be indicators of being stuck in an overall poor mode of responding, leading to poorer performance on these trials compared to trials triggered following regular ITIs.

#### Test 4. Time structure in the reaction time data

Kelly et al. (2001) found reduced temporal dependencies in a self-paced task compared to fixed-paced conditions. In our attempt to replicate this finding, both autocorrelations and power spectra were considered, following Wagenmakers et al. (2004). Autocorrelations measure the degree of dependency in a (reaction time) series with itself over time, by calculating the correlation between trial *n* and trial *n + k*, with *k* indicating the lag. Power spectra also measure temporal structures, but express this in frequencies – which allows for classification into different types of noise. Series with no temporal structures are called ‘white noise’, and are characterised by flat null autocorrelation functions as well as flat power spectra. It has been proposed that empirical data contains ‘pink noise’ or *1/f noise*, a mixture of strong short-term dependencies and slowly reducing long-term dependencies (Gilden, 2001; but see Farrell, Wagenmakers & Ratcliff, 2006; Wagenmakers et al., 2004), and is characterised by exponentially decreasing autocorrelation functions and power spectra with a slope around −1. Note that the power spectra can be mathematically derived from the autocorrelations.

If participants can use control to reduce their variability, the assumption is that they are accessing and mitigating against some internal state that fluctuates over time. Successful mitigation would result in reduced temporal dependencies in their RTs (i.e. closer to white noise) – reflected in reduced autocorrelations and flatter power spectra of the RTs in the self-paced condition. The temporal dependencies may instead be found in the self-paced ITIs (analysed in second section).

## Methods

### Participants

In total, 27 participants (20 female, 18-36 years, *M*_*age*_ = 25.6 years) with normal or corrected-to-normal vision were tested. Of them, 24 participated in the action-oriented task, and 27 participated in the perception-oriented task. We did not aim at a given Bayes Factor cut-off for participant recruitment, because we are considering multiple tests in parallel and it would have been very difficult to ensure that all of them reach such pre-determined Bayes Factor. Instead, the original aim was to recruit 24 participants, providing two repetitions of each block randomisation order. However, some participants dropped from the experiment following the perception-oriented task, leading us to recruit another 3 participants. We used Bayesian statistics on this first cohort and found the overall pattern of results sufficiently convincing (see Figure 9). Participants were paid £10/hour for participation. One participant in the action-oriented task and three in the perception-oriented task were excluded from analysis due to poor performance (see *Data preparation and analysis*). The study was approved by the local ethics committee.

### Materials

The stimuli were generated using MATLAB 8 (The MathWorks, Inc., Release 2016a) and Psychtoolbox-3 (Brainard, 1997; Pelli, 1997; Kleiner et al., 2007), using a Bits# Stimulus Processor video-graphic card (Cambridge Research Systems, Cambridge, UK) and a Viglen VIG80S PC (Viglen, Hertfordshire, UK), and were displayed on an hp p1230 monitor (Palo Alto, US) with a resolution of 1280 by 1024 and a refresh rate of 85Hz. Responses were recorded with a CB6 Push Button Response Box (Cambridge Research Systems, Cambridge, UK), which was connected to the Bits#. Participants were positioned in a chin- and head-rest, 92 cm away from the screen.

The experiment was shown on a grey background (55.8 cd/m^2^), featuring a fixation dot (112.1 cd/m^2^, .18°) or a fixation cross (112.1 cd/m^2^, .42°). Both tasks featured a vertically oriented Gabor patch as target (spatial frequency = 1.81 c/°, sigma = .26°). In the action-oriented task, the contrast of the target was always set at the maximum of 1. The perception-oriented task featured a low-contrast (difficult to detect) target that was adjusted to individual detection-thresholds of 75% accuracy and ranged between .021-.070 (*M* = .039, *SD* = .011).

### Design

Both tasks had four conditions: Self-Paced, Fixed, Replay and Shuffled replay. In the Self-Paced condition, participants started each new trial manually whenever they felt ready –they were given control over the ITI. In the Fixed condition, the median of the ITIs in the Self-Paced condition was used as ITI-length. The ITI was thus kept fixed throughout the trials while keeping the pace as similar as possible to the self-paced trials. In the Replay condition, the recorded ITIs from the Self-Paced condition were Replayed in the exact same order – thus controlling for the different ITI lengths without giving control to the participants – and in the Shuffled replay condition, the ITIs were Replayed in a different order – to allow for the different ITI lengths while removing any possible time structure between the ITIs.

### Procedure

The experiment consisted of four testing days of about an hour – two for each of the tasks (Figure 2). The first day of both tasks started with a training of 300 trials, followed by the Self-paced condition, and then one of the three control conditions (Fixed, Replay, or Shuffled replay). The remaining two conditions were administered on the next day. On each day, the testing session was preceded by three minutes of rest with eyes open, to provide a common baseline to all participants before starting the task.

**Figure 2.**
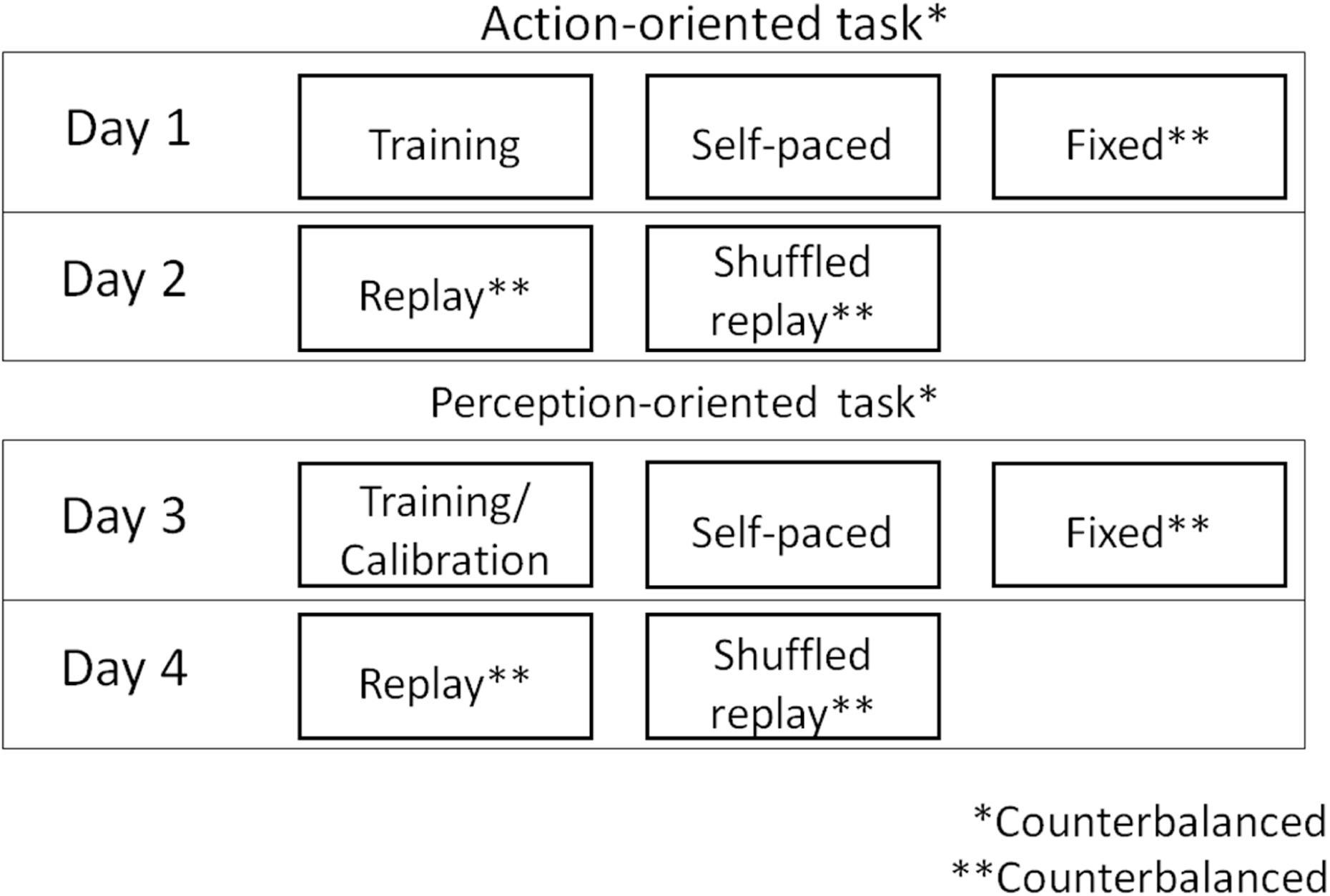
Structure of Experiment 2. Each task (action- and perception-oriented) took place over two days, with the order of the tasks being counterbalanced over participants. For both tasks, participants started with a training of 300 trials followed by the Self-paced condition, and finally one of the three control conditions (Fixed ITI, Replay, or Shuffled Replay, the order being counterbalanced over participants). During the next session, they would perform the other two conditions.

#### Main Experiment

Figure 3 illustrates the time course of each trial. Every trial started with a light grey screen with a fixation dot in the centre. Each condition consisted of 300 trials, with the first 30 being training trials. In the Self-paced condition, participants were instructed to press with the left and right index fingers at the same time whenever they felt ready for a new trial. They were told that they could wait as long as they wanted before continuing, but were discouraged from taking very long breaks. The time between fixation dot onset and double key press was recorded and subsequently used as ITI in the other conditions. Participants were unaware that their own self-paced ITIs would be used. After the button press, the dot was replaced by a fixation cross. In the three forced-paced conditions, the participant’s recorded self-paced ITIs (Replay, Shuffled replay) or median (Fixed) were used to determine the time between fixation dot and fixation cross. Next, 500ms after the cross onset, a target appeared either on the left or right of the cross. Participants were instructed to indicate with a button press which side the target appeared, using their left or right index fingers. After 200ms, the target disappeared, and after another 100ms, the fixation cross disappeared. Participants were then shown a blank screen until they responded.

**Figure 3.**
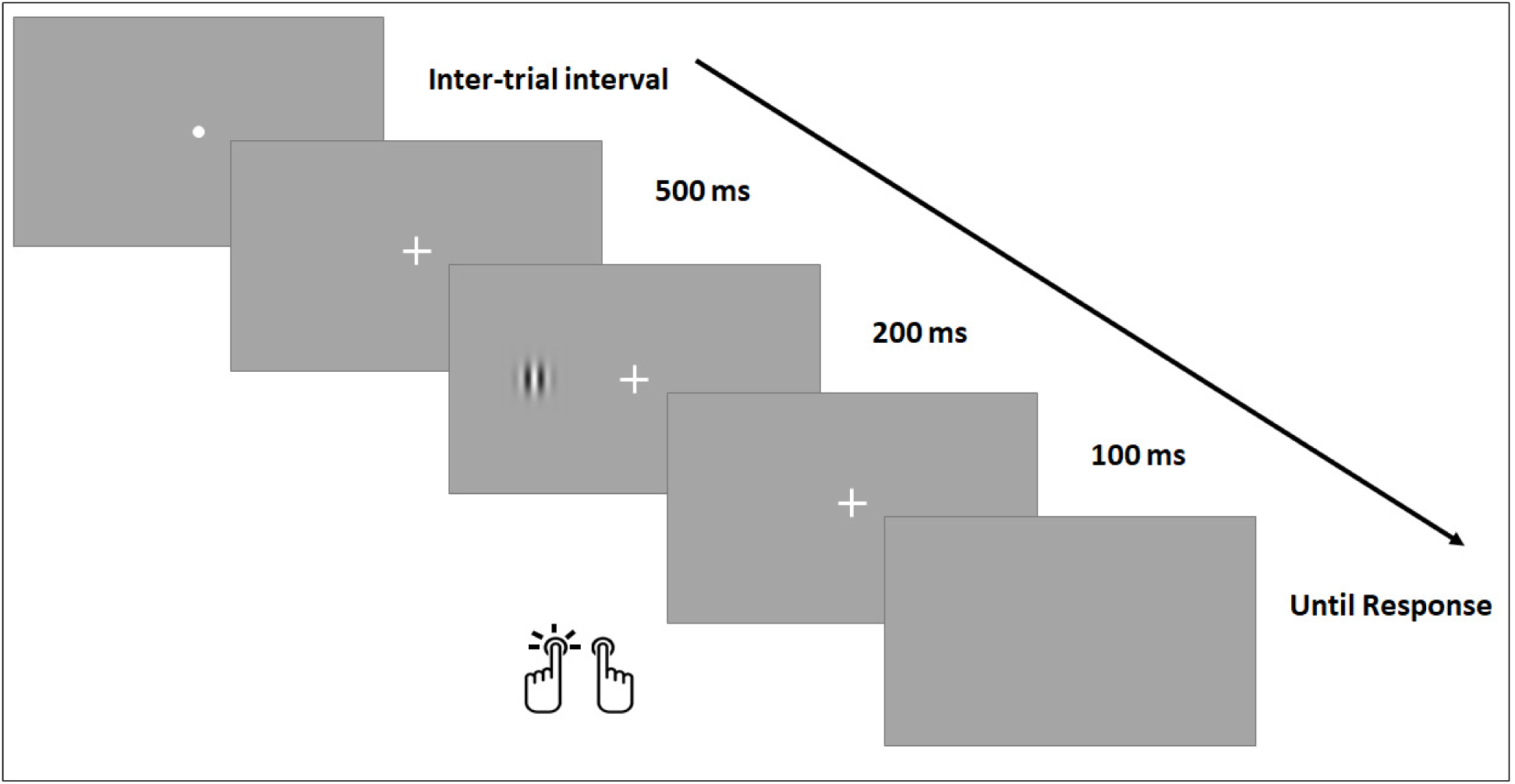
Example of one trial over time in Experiment 2. The length of the inter-trial interval was manipulated over conditions. After the ITI, the fixation dot was replaced with a fixation cross. After 500ms, the stimulus (Gabor patch) appeared either on the left or the right side of the screen for 200ms. The fixation cross disappeared 100ms later, and the screen remained empty until the participants responded either with their left or right index finger.

#### Training

Before the main experiment, each participant underwent a training using a fixed ITI of 1000 ms. After every 30 trials, participants were given feedback on their mean reaction time and accuracy. In the action-oriented task, participants were asked to be as fast as possible while avoiding errors, and in the perception-oriented task, they were asked to be as accurate as possible while avoiding producing too long RT. Again, the focus of the instructions was on good performance, and not on consistency. These instructions were repeated in the main experiment before each new condition. In the perception task, these trials were also used to determine the target contrast for each individual for the remainder of the task. The Psi method (Kontsevich & Tyler, 1999) was used to find the 75%-correct contrast detection threshold for each participant. Performance on training trials were excluded from all analyses.

## Results

### Test 1. Participants do not perform consistently better with control

Average RT across conditions and across participants ranged from 204 to 932 ms in the action-oriented task and from 271 to 2143 in the perception-oriented task. However, participants’ data were highly skewed, which had a large effect on the calculations of the mean (and variability) of the RT. Moreover, group distributions of mean RT and CVRT violated assumptions for normality. Therefore, RTs were log transformed. Because our hypotheses rely on the assumption that participants are motivated and able to perform the task, we first examined performance for each participant. One participant was excluded for both tasks due to below chance level performance on the training trials. Three participants in the perception-oriented task were excluded from analysis as more than 15% of their correct trials reflected poor performance in at least one of the conditions (both outliers, which log(RT) was higher than 3 standard deviations above the mean log(RT), and extreme RT (below 100 or above 1000 ms in action-oriented, and below 150 or above 1500 ms in perception-oriented task). As these participants tended to perform poorly in all conditions, this should not bias either hypothesis.

Examining the unfiltered data of the remaining participants, average RT across conditions ranged from 204 to 592 ms in the action-oriented-task and from 271 to 1550 ms in the perception-oriented task. Mean accuracy scores were calculated for each participant and for each condition. Mean reaction times and standard deviations were calculated on the logged values of the correct trials.

For both tasks, Test 1 involved paired Bayesian t-tests conducted on: 1) reaction time, 2) percentage of errors, and 3) CVRT, to test if the self-paced condition differed from any of the three control conditions. Because Hypothesis 1 is specifically based on *better performance* in the self-paced compared to the other three conditions, the t-tests were conducted one-sided.

Figure 4 compares the Self-paced condition to each of the forced-paced conditions on individual measures of performance (RT and percentage of errors) and intra-individual variability (CVRT). Table 4 shows the corresponding Bayes’ Factors. Altogether, we did not find any consistent benefit of the Self-paced condition over the forced-paced conditions, and evidence overall favoured *H2* over *H1*. Below the results are described in more detail.

**Table 4.** Statistical outcomes for Test 1 – Does control improve performance and reduce variability? Shown are the Bayes’ Factors for H1 over H2 for each comparison on RT, % errors and CVRT on both the action-oriented and the perception-oriented task. T-tests were conducted one-sided by contrasting the Self-Paced (SP) to each of the 3 forced-paced conditions, Fixed-paced (F), Replay (R) and Shuffled Replay (SR).

| BF<sub>12</sub> | Action-oriented |  |  | Perception-oriented |  |  |
| --- | --- | --- | --- | --- | --- | --- |
| Comparison | RT | % Errors | CVRT | RT | % Errors | CVRT |
| SP < F | .58 | .39 | .39 | .06 | .33 | 7.70 |
| SP < R | 51.07 | .06 | 1.07 | .01 | .57 | 123.97 |
| SP < SR | 333.62 | .05 | .48 | .13 | .53 | 2045.39 |

**Figure 4.**
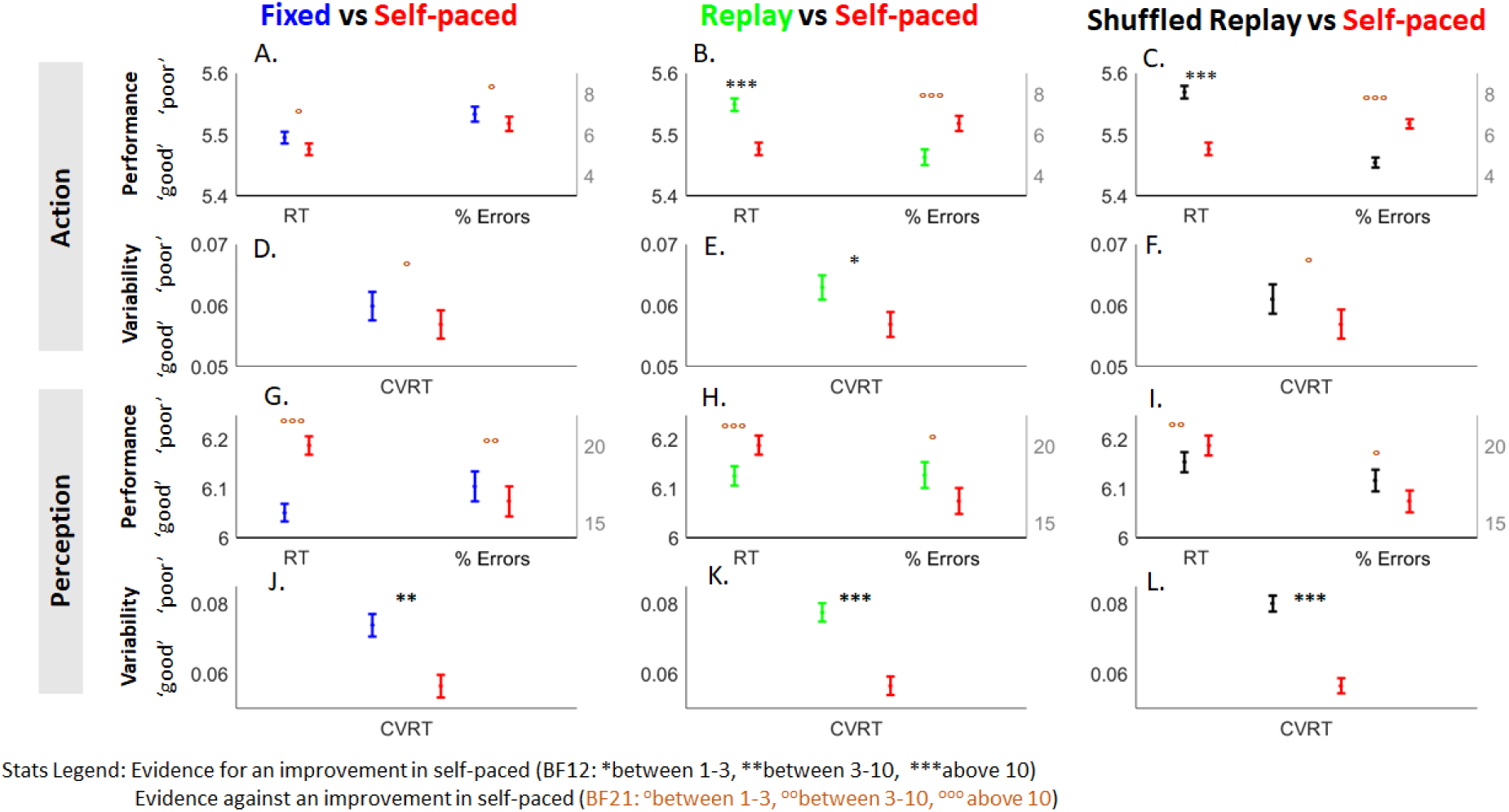
Mean log RT, percentage of errors, and CVRT in the Self-paced condition compared to each of the three forced-paced conditions: A) Fixed ITI (blue), B) Replay (green) and C) Shuffled Replay (black). Black stars indicate evidence for an improvement in the Self-paced condition (consistent with H1), while orange circles indicate evidence against an improvement in the Self-paced condition (against H1). Top and Bottom panels show results of the action- and perception-oriented task. Error bars show the within-subject standard error across conditions. Increasing scores on the Y-axes show decreasing performance in a single measure (lower speed, accuracy or consistency), but the measures should be interpreted in relation to each other.

#### Performance

Altogether, none of the comparisons in both tasks revealed any clear benefit of the control on performance. In the action-oriented task, the comparisons with the Fixed condition (Figure 4A) were both in the indeterminate range (BF between 1 and 3), while the comparison with the Replay and Shuffled replay conditions showed that participants were on average faster in the self-paced condition (providing strong evidence for *H1*), but also made more errors (providing strong evidence against *H1*, Figure 4B-C). This pattern is actually suggestive of an adjustment in speed-accuracy strategy, probably in response to the target onset being predictable (versus unpredictable in the Replay and Shuffled Replay conditions), rather than the improvement in performance expected under *H1*. This interpretation is supported by modelling using the EZ-Diffusion Model (see Test 1b). In the perception-oriented task (Figure 4G-I), all six comparisons favoured *H2* (BF21 ranging from ∼1.8-100), though two were in the indeterminate range.

#### Variability

In the action-oriented task (Figure 4D-F), all the comparisons were in the indeterminate range, and therefore not showing any clear benefit of control on CVRT. In the perception-oriented task, all the comparisons showed moderate to strong evidence for *H1*, i.e. lower CVRT in the Self-paced condition compared to the forced-paced conditions. It is noteworthy that such reduced intra-individual variability was not accompanied by a reduction in mean RT (in fact, mean reaction time was highest in the Self-paced condition). One interpretation could be that participants made less anticipatory responses in the Self-paced condition, possibly due to the additional button press to initiate the trial. In this case, this reduction in CVRT would not be interpreted as an improvement in performance, but rather as an indication that participants are behaving differently in the self-paced condition. It should also be noted that this decrease in anticipatory responses did not lead to an increased accuracy, even though anticipations are characterised by accuracy scores at chance level (suggesting that a reduction in them would increase overall accuracy). Test 1b and Test 2 below address this in more detail.

### Test 1b. EZ-model suggests strategy-adjustments, not performance improvement

Drift rate, boundary separation, and non-decision time parameters were calculated for both tasks on each condition. Because the estimations are sensitive to outliers, extreme high RT (1000 ms for the action-oriented and 1500 ms for the perception-oriented task) were excluded before calculating the parameters. Next, Bayesian Paired t-tests were performed on i) drift rate (specifically testing one-sided for *increased* drift rate in the self-paced condition compared to the other three conditions, which may reflect improved performance), ii) non-decision times (specifically testing one-sided for *decreased* non-decision times in the self-paced condition compared to the other three conditions), and iii) boundary separation (specifically testing two-sided for any difference between the conditions, reflecting changes in response strategies).

Figure 5 shows the means of drift rate (v), non-decision times (T_er_), and boundary separation (α) as calculated by the EZ-Diffusion model in the Self-paced condition compared to each of the forced-paced conditions, with corresponding Bayes Factors shown in Table 5. The first two parameters may reflect differences in performance (with good performance being indicated by *higher* drift rate and *lower* non-decision times), while boundary separation indicated differences in speed-accuracy trade-off (with higher values indicating a more cautious strategy).

**Table 5.** Bayes’ Factors contrasting the EZ-diffusion parameters between the Self-paced and each Forced-paced condition (v: drift rate; T_er_: non-decision time; α: boundary separation). Same conventions as Table 4.

|  | Action-oriented |  |  | Perception-oriented |  |  |
| --- | --- | --- | --- | --- | --- | --- |
| Comparison | $v^*$ | $T_{er}^{**}$ | $\alpha^{***}$ | $v^*$ | $T_{er}^{**}$ | $\alpha^{***}$ |
| SP - F | .50 | .46 | .28 | .41 | .23 | .06 |
| SP - R | .20 | .40 | 2.99 | .95 | .26 | .06 |
| SP - SR | .20 | 1.37 | 9.74 | .98 | .25 | .07 |

\*Tested for higher drift rates in the self-paced condition than the other conditions.
\*\*Tested for lower non-decision times in the self-paced condition than the other
conditions.
\*\*\*Tested for no difference between the conditions.

**Figure 5.**
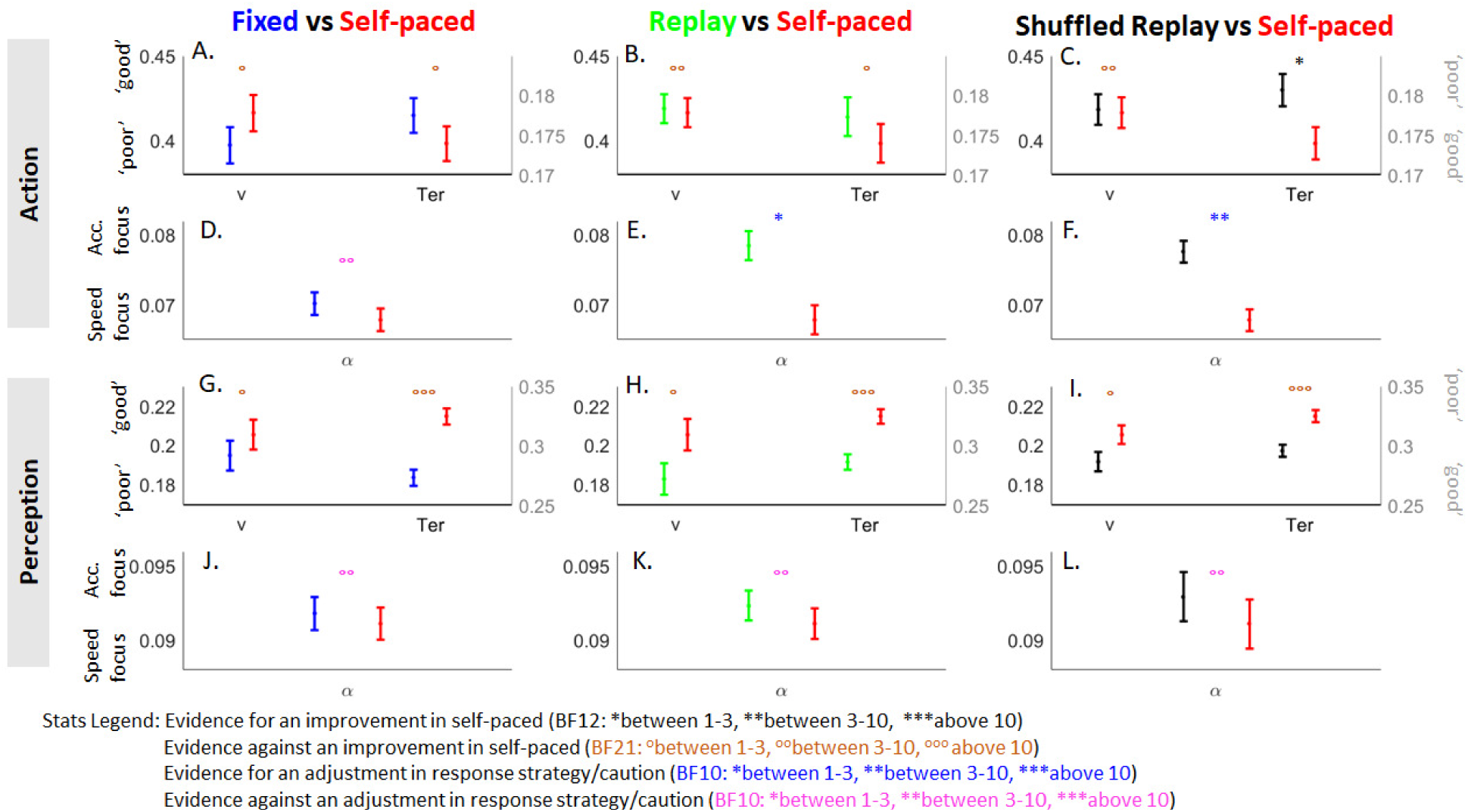
Averages of the EZ-diffusion parameters on the Self-paced (SP), Fixed (F), Replay (R), and Shuffled Replay (SR) conditions. Error bars show the within-subject standard error.

Altogether, in the action-oriented task, the comparisons suggest that the differences between conditions are caused by adjustments in speed-accuracy trade-off. These adjustments seem dependent on predictability of target onset rather than on control. This supports the conclusion that there is no benefit of control on performance – supporting *H2* over *H1*. For the perception-oriented task, differences are best explained by an increase in non-decision times, and thus, a decrease in performance. Again, this supports *H2* rather than *H1*. Below the results are described in more detail.

#### Performance

In the action-oriented task, there was no consistent improvement in the Self-paced condition (Figure 5A-C). Out of the six comparisons, only one favoured *H1* (reduced non-decision times in Self-paced compared to Shuffled Replay), but the evidence was indeterminate (BF close to 1). In the perception-oriented task, there was strong evidence against a decrease in non-decision times in the Self-paced condition compared to each of the forced-paced conditions (Figure 5G-I). In fact, non-decision times were higher in the Self-paced condition, with no evidence for increases in drift rate. This clearly suggests that processing of information did not improve in the Self-paced condition compared to the forced-paced conditions, but rather, that sensory or motor processes took longer (see Test 2 for complementary evidence).

#### Speed-accuracy strategies

In the action-oriented task, indeed, boundary separation in the Self-paced was lower compared to the Replay and Shuffled replay condition (Figure 5B-C). High boundary separation values indicate that a lot of information needs to be gathered before one option can win (thus taking longer on the decision process, but with fewer chances of errors), while low values indicate that less information needs to be gathered before one option can win (leading to shorter RT, but reduced accuracy). Further testing showed that boundary separation was also lower in the Fixed condition compared to Replay and Shuffled replay (BF of 11.1 and 13.8 respectively), confirming that participants were less cautious when the target onset was predictable. In the perception-oriented task, there was strong evidence against a change in boundary separation (figure 5J-L). As such, it seems there are no differences in speed-accuracy strategies in the perception-oriented task between the Self-paced and the forced-paced conditions.

### Test 2. Differences in extreme RTs

The amount of extreme reaction times (including trials defined as outliers above) in each condition was calculated for each of the participants. As a lower-bound cut-off, trials were counted if the RT was below 150 ms or below 200 ms for the action-oriented task and the perception-oriented task respectively. Trials below these cut-offs showed chance performance (i.e. an average accuracy of 50%) and as such reflect anticipations. For the upper-bound cut-off, trials were counted if the RT was above 500 ms or 1000 ms for the action-oriented and perception-oriented task respectively. Bayesian Paired one-sided t-tests were conducted, testing if the number of extreme reaction times was lower in the Self-paced condition compared to each of the fixed-paced conditions.

In the action-oriented task, anticipations were not less frequent is the Self-paced condition compared to each of the three fixed-paced conditions (see Figure 6A-C for means and Bayes’ Factors), providing moderate to strong evidence for *H2* over *H1*. The number of high reaction times were more mixed: While there was moderate evidence against *H1* in the comparison with the Fixed-paced condition, the BF for Replay and Shuffled Replay were both close to 1.

**Figure 6.**
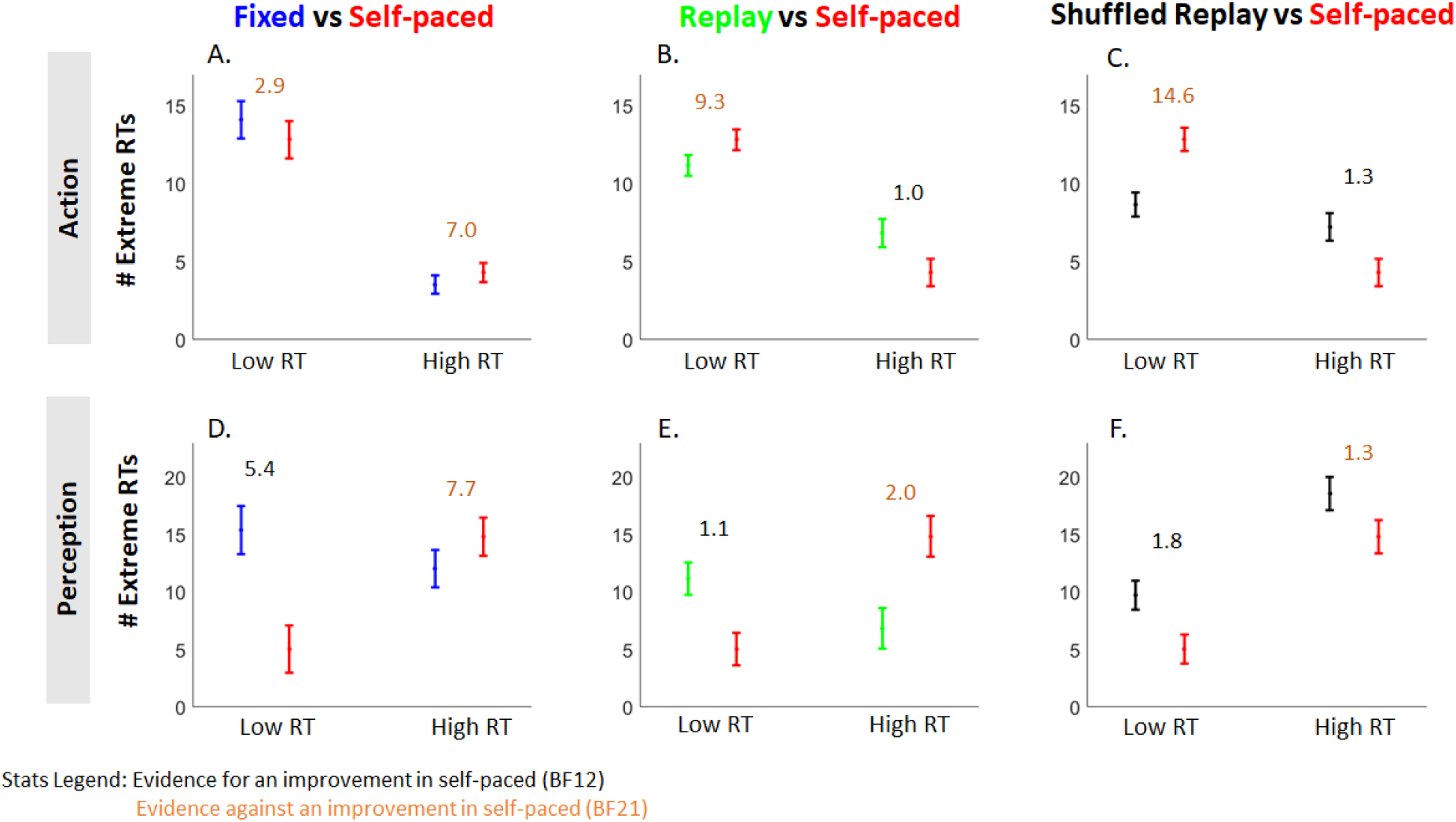
Number of extreme reaction times averaged across participants. Same conventions as Figure 4.

In the perception-oriented task (see Figure 6D-F), support for *H1* was found only for anticipations, while the high RTs favoured *H2* overall. This reduction of the very short reaction times in the Self-paced condition was consistent with the overall higher mean RT compared to all the forced-paced conditions – bringing support to the interpretation from Test 1. One possibility is that this is due to the interference of the additional button press in the Self-paced condition. This interpretation is consistent with our modelling using the EZ-Diffusion model, which suggested that only non-decision times were higher in this condition.

### Test 3. Longer ITIs lead to poorer performance, not better

ITI-distributions were calculated by taking the time between the response on trial *n-1* and the self-paced ITI-press on trial *n*. To define typical and longer ITIs, the RT distribution from the Fixed condition was used as a reference: For both tasks, the 95^th^ percentile of the RT-distribution was calculated for each participant as cut-off (see Figure 6A for an example). Self-paced ITIs below this cut-off were classified as ‘regular’, and may reflect as fast as possible responses to the fixation dot indicating one can start a new trial – thus resembling regular RT. ITIs above the cut-off were classified as ‘long’, and may reflect times in which participant felt they needed to wait longer before feeling ready to continue.

Mean error scores, mean reaction times, and standard deviations were calculated for trials following on from regular ITIs as well as for trials following long ITIs. Because there was a lot of variation in the number of regular and long trials between participants, ten trials were randomly selected 10000 times and the mean accuracy, reaction time, and CVRT over these 10000 iterations were calculated. Subsequently, Bayesian paired one-sided t-tests were conducted on these means to see if performance improved following long ITIs. In the action-oriented task, three participants were excluded from analysis because they had less than ten trials with regular self-paced ITIs.

For both tasks, evidence was found against an improvement in RT, variability or accuracy –providing moderate to strong evidence against *H1* (Figure 7B). When testing in the opposite direction (long ITIs lead to worse performance), it was found that RT and variability (but not accuracy) were clearly worse following long ITIs than those following regular ITIs (BFs of 13.23 and 15.84 for the action-oriented task, and 155.56 and 77.26 for the perception-oriented task respectively).

**Figure 7.**
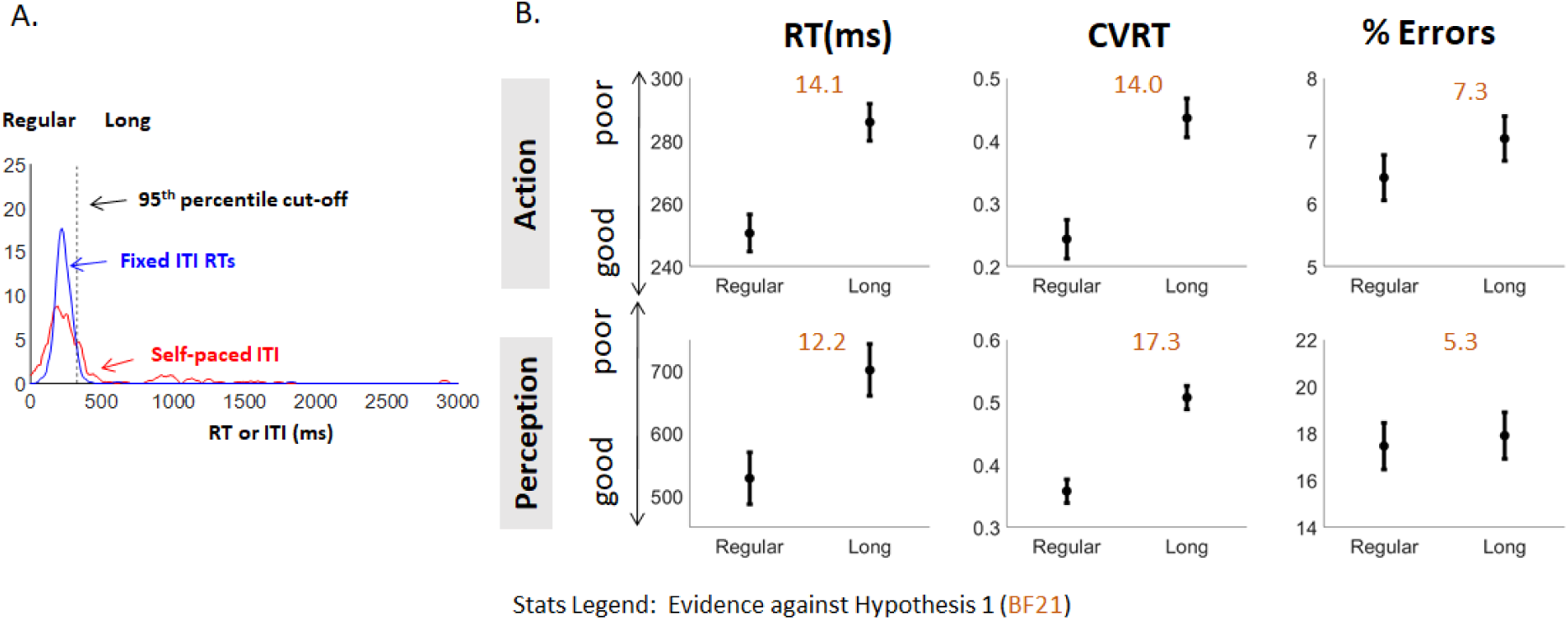
Detrimental effect of long self-paced ITIs on performance and variability. **A)** Example from one participant of regular and long ITI-trials. Shown are the smoothed distribution of the self-paced ITIs (in red) and the distribution of the RT of the Fixed ITI condition (in blue). For each participant, the 95^th^ percentile of the Fixed ITI RT distribution was calculated as a cut-off (black dotted line). Self-paced ITIs above the cut-off were deemed ‘long’, while ITIs below the cut-off were deemed ‘regular’. ***B)*** Mean RT, CVRT, and % errors were calculated in the Self-paced condition for trials following regular and long ITIs. Orange Bayes’ Factors indicate the strength of evidence against H1. None of the comparisons were in favour of H1. Error bars show the within-subject standard error.

In conclusion, long self-paced ITIs did not lead to an improvement in performance or a reduction in variability. Instead, these breaks were associated with subsequent lower performance and higher variability. The co-occurring long ITIs and longer reaction times suggest the same fluctuating internal states affect both measures. To confirm this, correlation coefficients between ITI and RT on each trial were also performed. For both tasks, correlation coefficients were positive overall on the group (BF for one sample t-tests 17.1 and 18) – suggesting that short ITIs are typically followed by short RTs, and long ITIs by long RTs. This could reflect similar temporal dependencies as in typical RT series on consecutive trials.

### Test 4. Control does not reduce temporal dependencies in RT series

The autocorrelations in the reaction time data were calculated separately for each participant and condition. Furthermore, the power spectrum was calculated over each reaction time series in R 3.3.2 (R Core Team, 2016), following Wagenmakers et al. (2004). Although Wagenmakers et al. (2004) showed that the power spectrum for white noise of variance 1 is flat and null, this is not the case for white noise with the same variance as our experimental data, nor for series obtained from randomly shuffling our data. Therefore, to correct for the power spectrum expected in our time series irrespective of any temporal dependency (our null hypothesis), the power spectrum was calculated 100 times on the randomly shuffled reaction time data, and the mean of these 100 spectra was subtracted from the unshuffled power spectrum. As such, the difference of these spectra reflects the time structure in the reaction time data. These difference-spectra were calculated separately for each participant and each condition.

Next, a linear regression line was fitted on the log of each power spectrum (still following Wagenmakers et al., 2004). Paired Bayesian t-tests were then conducted on the autocorrelations at the first lag and on the spectral slopes – to test if the self-paced condition differed from any of the three forced-paced conditions. Again, because *H1* is based specifically on a decrease in temporal dependency (and thus a flatter slope), t-tests were conducted one-sided.

First, we checked that our RT and ITI series actually showed clear temporal structure. As there was consistent evidence for dependencies across the two measures (See Supplementary Table 1), we carried on with contrasting these temporal dependencies across conditions.

Figure 8 shows the mean autocorrelation functions and power spectra. Table 6 shows the Bayes’ Factors associated with comparing each forced-paced condition to the Self-paced condition. Across both tasks, all comparisons except one provided evidence for *H2* over *H1* (showing no decrease in temporal dependencies in the Self-paced condition). The only exception was the decreased spectral slope compared to the Fixed condition in the action-oriented task, which showed only indeterminate evidence for *H1*.

**Table 6.** Bayes’ Factors for Test 4, comparing temporal dependencies in the Self-paced versus each forced-paced condition, as reflected in the first point of the autocorrelation (AC) and the fitted slopes on the spectral power. Same conventions as Table 4 and 5.

|  | Action-oriented |  | Perception-oriented |  |
| --- | --- | --- | --- | --- |
| Comparison | AC | slope | AC | slope |
| SP < F | .74 | 2.05 | .14 | .30 |
| SP < R | .51 | .82 | .10 | .19 |
| SP < SR | .13 | .14 | .06 | .09 |

**Figure 8.**
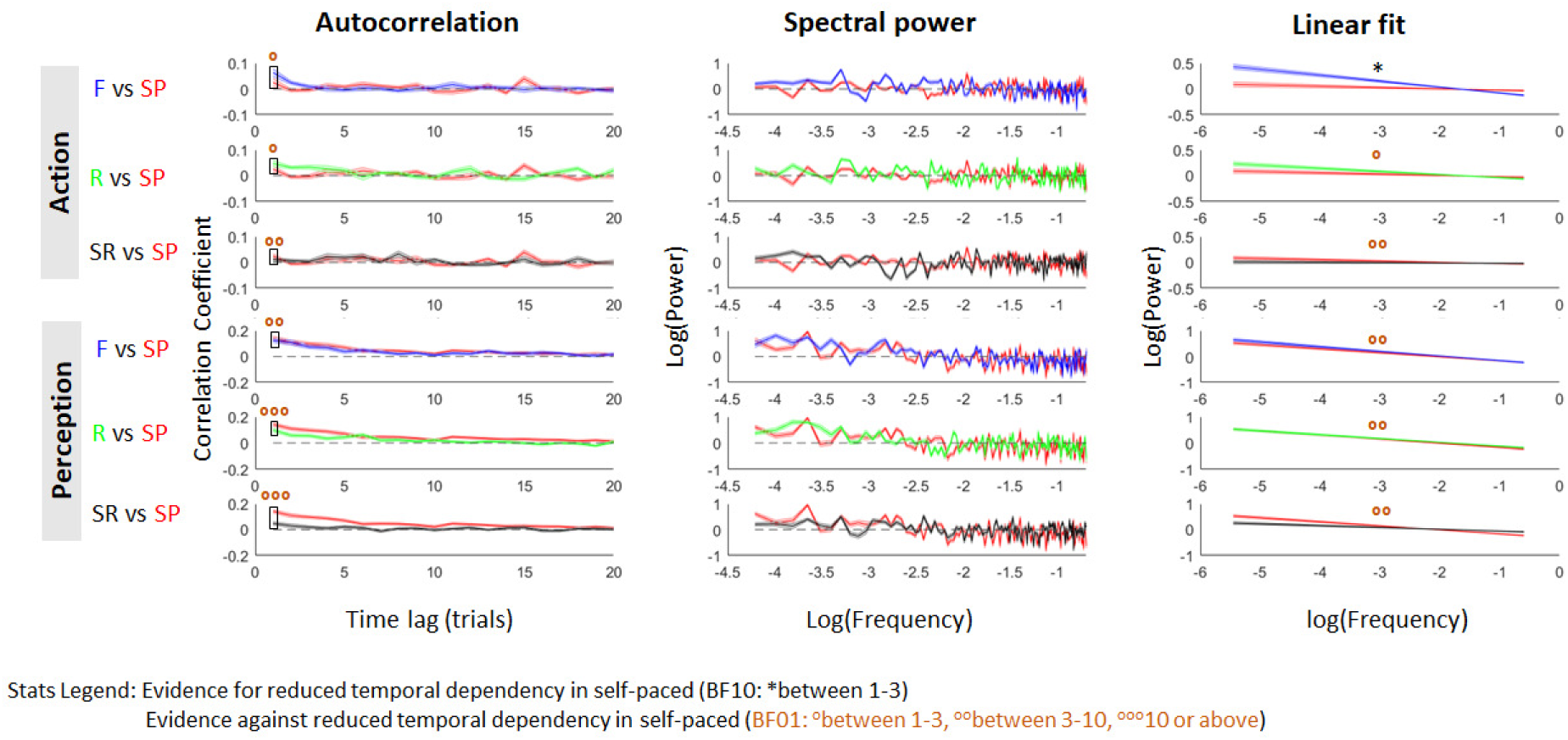
Autocorrelation and spectral power and corresponding linear fit over the spectral power averaged across participants (same conventions as in Figures 4 and 5) for the RT, comparing the Self-paced with each of the forced-paced conditions.

**Figure 9.**
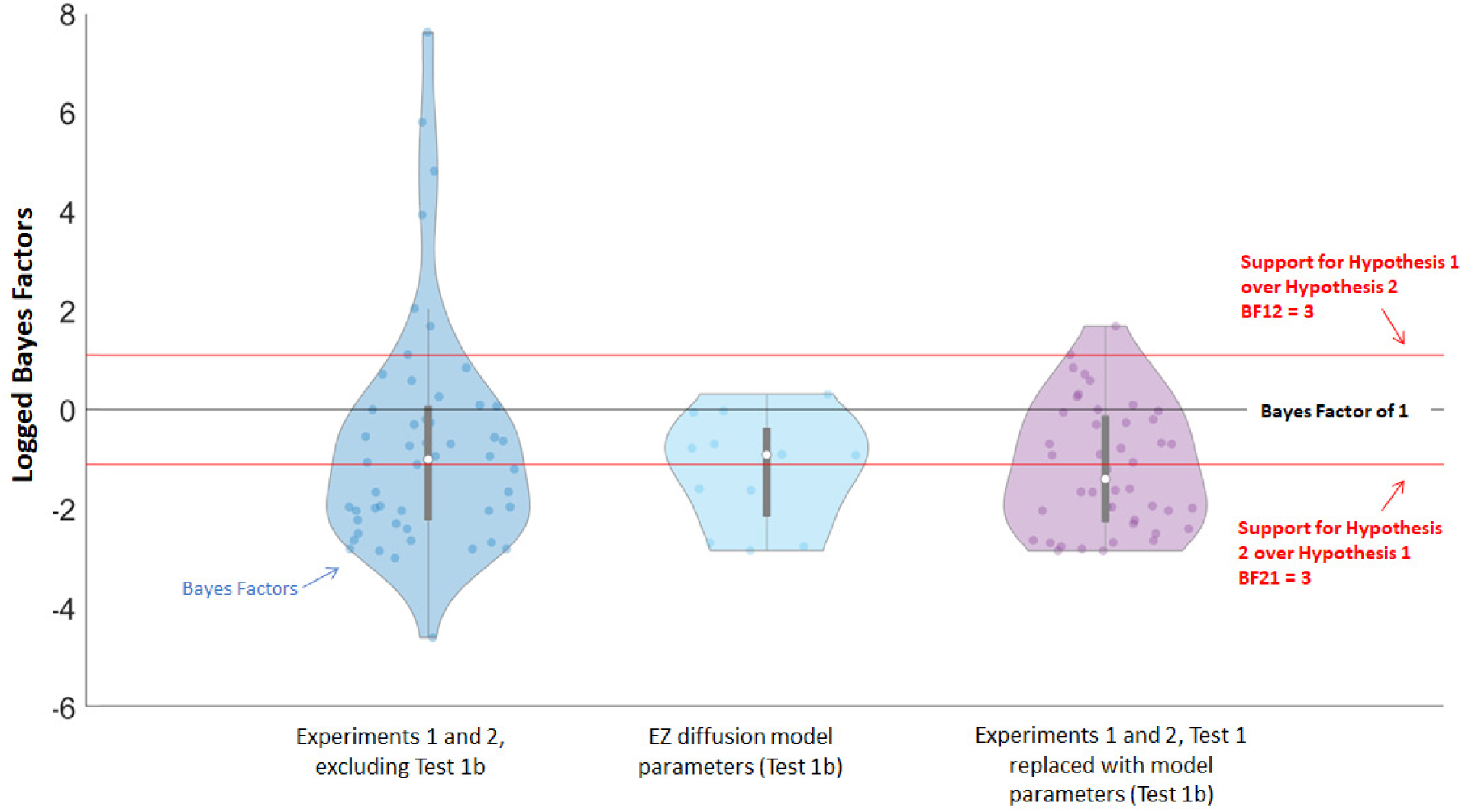
Distribution of logged Bayes Factors from the statistical tests that compared Hypothesis 1 to Hypothesis 2, with each coloured dot representing one Bayes Factor, and each white dot representing the median of that distribution. Dots above the black line reflect higher support for Hypothesis 1, while dots below the black line reflect higher support for Hypothesis 2. The most left distribution (dark blue) encompasses the Bayes Factors from Experiments 1 and 2 (Test 1-4, excluding the EZ-model comparisons from Test 1b). In the right distribution (purple), the comparisons of RT, CVRT, and percentage correct (Test 1) have been replaced by the comparisons of the modelling on drift rate and non-decision times (Test 1b – shown separately in the middle graph). Overall, the distributions show our results favour Hypothesis 2 over Hypothesis 1.

Overall, our results suggest that control over trial initiation does not affect temporal dependencies. The only exception was found in the action-oriented task, when comparing the Self-paced condition to the Fixed condition. Although the evidence was in the indeterminate range and should thus be interpreted with high caution, this seems in line with previous research, also using an action-oriented task, which found lower temporal dependency in a self-paced and irregular condition compared to fixed-paced and regular conditions (Kelly et al., 2001). Thus, if the effect exists, it is more likely to be due to the difference in regularity of target onset rather than self-paced control per se.

### No training effects in the Self-paced condition

Because the three fixed-paced conditions depended upon participants’ own self-paced ITIs, the Self-paced condition always had to come first – making full counterbalancing impossible. While the potential effects of this are not straightforward, we conducted an extra analysis to test if participants were still learning the task in the Self-paced condition even after the training block. For each condition, the mean RT and accuracy were calculated for each participant on: 1) the first 30 trials (excluding the first trial), and 2) the rest of the trials. A Bayesian paired t-test was conducted to test if participants performed worse on the first set of trials than on the rest of the experiment (reflecting training effects).

No differences were found in either RT or accuracy in either task between the first 30 trials and the remaining trials in the Self-paced condition, (BF01 = 14.3, 19.1, 3.2 and 14.9 for RT in action-oriented task, accuracy in action-oriented task, RT in perception-oriented task and accuracy in perception-oriented task respectively). It is thus unlikely that any outcome of the analyses from Experiment 2 could be ascribed to condition orders.

## Interim discussion 2

Figure 9 summarises all the Bayes Factors of the statistical comparisons between *H1* and *H2* (combining Experiment 1 and 2) with violin plots – distribution plots that show the entire range of Bayes Factors (y-axis) with horizontal thickness indicating density. Note that the Bayes Factors are logged for graphical purposes. The most-left (dark-blue) violin represents Experiment 1 plus Tests 1-4 of Experiment 2, excluding the EZ-parameter comparisons from Test 1b – showing an overall bias towards *H2*. While there are some BF that highly favour *H1*, these relate to comparisons that likely represent differences in speed-accuracy trade-offs, and do not reflect actual improvements in performance.

In the most-right violin (purple), the comparisons on RT, CVRT, and percentage correct have therefore been replaced by the comparisons between the parameters of the EZ-Diffusion model that relate to performance (drift rate and non-decision times from Test 1b, also seen separately in the middle violin). The comparisons on boundary separation are not included because they do not favour either hypothesis by default. Again, the overall results favour *H2*, showing evidence *against* a benefit of control.

## Characterising the self-paced ITIs in the computer-based tasks

### Rationale

The results from Experiment 1 and Experiment 2 suggest that participants do not benefit from having control. In the remainder of the article, we focus on *how* participants used the control. Because Experiment 2 allows for the recording of the self-paced ITIs, it provides an opportunity to examine these ITIs in more detail – to see what potential strategies participants may have used in handling the control they were given. Although the control did not benefit participants, their ITIs may still show characteristics that diverge from regular RT characteristics. To get more insight into these strategies, we examined three different measures in the self-paced ITIs: Variability, temporal dependencies, and post-error slowing.

#### Variability in the ITI

If participants use the control in the self-paced condition and do not continue to the next trial when they do not feel ready, one would expect the distributions of the self-paced ITIs to be different from typical responses to a stimulus. Specifically, if participants make use of the control, they should show a mixture of shorter and longer ITIs – which subsequently leads to high variability. On the other hand, if participants just start the trials as soon as the stimulus inviting them to do so appears, their ITIs should resemble simple RTs to a single salient and predictable stimulus onset (the fixation cross). We did not have such data from our participants but the Fixed condition from the action task offered the closest comparison. If participants were just eager to carry on through the task as quickly as possible, the coefficient of variation of their ITIs (CVITI) across both tasks should be similar to the CVRT from the Fixed condition in the action task, or even smaller, because it is a one-alternative decision, while the RT is based on a two-alternative decision.

#### Temporal dependency in the ITI

As mentioned in the introduction, we expect the self-paced ITIs to show temporal dependencies. Because participants were instructed to wait for every trial until they felt ready for it, their ITIs may show higher temporal dependencies than typical RTs – possibly reflecting stronger coupling to fluctuating internal states than stimulus-driven responses (the trial itself), even if these attempts did not result in better performance. To examine this, the autocorrelations and power spectra were calculated for the self-paced it is. Again, for both tasks, autocorrelations and fitted lines were compared against the Fixed condition of the action-oriented condition.

#### Post-error slowing in the ITI

There is a large literature showing that people are able to slow down when they see or are explicitly told that they made an error (post-error slowing; Rabbitt, 1966) – seemingly because of an adjustment of response caution (Dutilh et al., 2012). When participants are making an error, they are faced with objective information that their performance-relevant internal state – and thus their decision to continue to the next trial – was suboptimal. If participants were able to make maximum use of the control based on their inner states, they could have prevented these errors from happening altogether, especially in the action task, which is very easy. However, since they were not able to use the control in this manner, they may instead slow down afterwards – resulting in post-error slowing in the ITI. This may at least indicate that our participants cared enough to adjust their behaviour in response to poor performance, even if this was ineffective in boosting their performance.

## Results

### ITIs show higher variability than RT

Mean ITI ranged from 243 to 1742 in the action-oriented task and from 298 to 2605 in the perception-oriented task. Similarly, to the RT data, the ITI data was log transformed as a first step, to correct for the high skew of the distributions. Figure 10A shows the distributions of the CVITI for both tasks compared with the CVRT of the Fixed condition of the action-oriented task, with accompanying Bayes Factors for the associated Paired one-sided t-tests. On both tasks, we found extreme evidence that the CVITI was much higher than the CVRT – showing that the self-paced ITIs are more variable than would be expected if they were just response times to a stimulus. This suggests that participants were using the ITI in some manner, but this did not help them to improve their subsequent performance.

**Figure 10.**
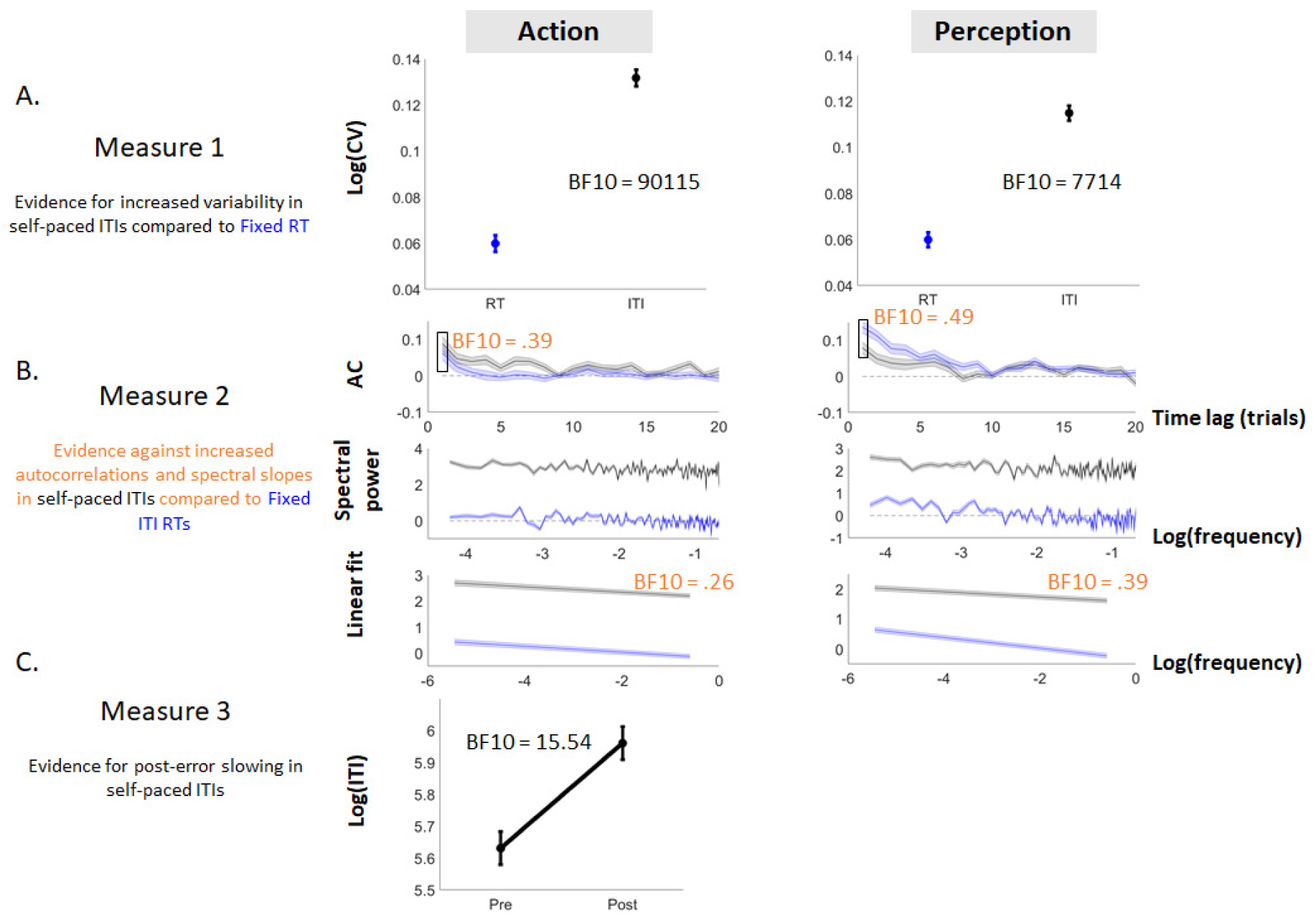
Evidence across the three measures of Part 2. Measure 1 reflects the coefficient of variance for the log of the ITI (CVITI) on both tasks, compared to the coefficient of variance for the log of the RT (CVRT) of the Fixed condition on the action-oriented task. Measure 2 reflects the temporal dependency of the ITI, as measured by the autocorrelation and the fitted slope on the spectral power, compared to that of the RT of the Fixed condition in the action-oriented task. The ITI showed much higher variability than the RT, but did not show higher autocorrelations or steeper slopes. Measure 3 reflects post-error slowing found in the ITIs. Data points show the logged average self-paced ITIs in the action-oriented task before (pre) and after (post) an error, indicating that participants slow down in their ITI after an incorrect trial. Error bars on all panels show the within-subject standard error.

### ITIs may show some higher temporal dependencies than RT

For both tasks, autocorrelations and power spectra plus their fit lines were calculated on the ITIs for each participant, using the same procedure as in Test 2 in above. Bayesian Paired one-sided t-tests were conducted on the autocorrelations at lag one and on the spectral slopes – to test if the temporal dependency was higher in the ITI compared to the RT of each condition. Similarly to Test 4 above, we first confirmed that the ITI actually contained temporal dependencies (see Supplementary Table 1). As we found evidence for this on both tasks, we then carried on with comparing the ITI to the RT.

Figure 10B shows the mean autocorrelation functions and power spectra of the ITIs from both tasks, compared to the RT of the Fixed condition of the action-oriented task. On both tasks, there was no evidence for higher temporal dependencies in the ITI compared to the RT.

### Post-error slowing in the self-paced ITIs

Post-error slowing in the self-paced ITIs was calculated using the method of Dutilh et al. (2012). To avoid unstable means due to a low number of observations, participants who made less than ten errors were excluded. For the remaining seventeen participants, mean ITIs were calculated on the logged ITIs before and after each error. Bayesian paired one-sided t-tests were performed to test if post-error ITIs were on average slower than pre-error ITIs. Because participants were not given feedback throughout the main tasks, post-error slowing was only calculated for the action-oriented task, in which participants typically know when they have made an error – as opposed to the perception-oriented task, in which participants are often unsure of the correct answer.

Participants were on average 215 ms slower in their ITI after making an error (Figure 10C – analysis run on logged values) compared to just before making this error. Such difference could have two possible origins though: 1) errors may lead to ITIs larger than average on the next trial, indicative that participants have adjusted their ITI as a consequence of the error (actual post-error slowing), or 2) errors could be typically preceded by shorter ITIs and followed by regular ITIs, simply reflecting a regression to the mean. Comparing the mean pre-and post-error ITI with the overall mean ITI shows clear support for option 1 (Bayesian one-sided paired t-test on logged values, BF10_post>mean_ = 8.86) and not for option 2 (BF10_pre<mean_ = 1.14).

It therefore seems that our participants were able to adjust their behaviour in response to *objective* evidence that their performance was poor (see section *Motivation* in the Discussion for more discussion on this), which is interesting for two reasons. First, this contrasts with their inability to adjust their ITIs to prevent errors from occurring, i.e. presumably in response to internally-driven information that they are in a state detrimental to performance. Second, it suggests they were sufficiently motivated to act upon their performance, which is a prerequisite for the control manipulation to be relevant.

The presence of post-error slowing could suggest that participants were able and willing to make some use of the control when faced with objective information on their performance. For this to lead to improved performance though, post-error slowing on trial *n* should also result in improved performance (i.e. lower RT and higher accuracy) on trial *n+1*, as focus has suddenly gone up. Unfortunately, neither of the current tasks are suited to examine this prediction, because post-error improvements in accuracy cannot be estimated properly (Danielmeier & Ullsperger, 2011): The action-oriented task has too few errors, leading to an unreliable estimate, and the perception-oriented task contains errors of which participants are not aware, which should not lead to subsequent post-error adjustments.

However, while the ITI could potentially absorb the slowing typically seen in RT, this may not necessarily lead to improved focus. If anything, the results of Test 3 above suggest that slowing down does not necessarily improve subsequent performance. Indeed, previous literature has shown that, while post-error slowing is often seen as a strategic adjustment aimed at improving subsequent performance, post-error slowing and post-error improvement in accuracy are not necessarily found together (see Danielmeier & Ullsperger, 2011 for a review). One possible reason could be that post-error slowing partly reflects an automatic response to rare events, similar to startling in the rodent literature (Wessel and Aron, 2017), rather than a purely strategic adjustment. The observed post-error slowing in the ITIs may as such reflect a mixture of automatic responses and top-down strategies to try to refocus on the task.

## General Discussion

### No improved performance or reduced variability with control

Assuming that task performance is under the influence of some internal states varying over time, we aimed to test whether people have direct access to these internal states and can use this information to improve task performance. We gave participants control over the timing of three behavioural tasks and compared their performance with conditions without such control. In all three tasks, we found that participants did not perform better when provided with control (see Figure 9 for an overview), even when questionnaires indicated high intrinsic motivation to perform the task. Furthermore, when participants took longer delays during the task, this was associated with poorer, not better, subsequent performance and increased variability. Control also did not affect temporal structures in the reaction times

When examining the time taken to move from one trial to the next in the self-paced condition (ITI), it is clear that participants do not simply rush through the task as quickly as possible. Rather, their ITIs are slower and show much higher variability than their speeded RT, as well as clear evidence of post-error slowing. This suggests participants use the control in some manner, but this did not help them improve their performance.

### Access to internal state: either limited or not directly useful

Overall, our results show that participants were not able to use the control to improve their performance and reduce their variability – suggesting that if people have some access to their performance-relevant inner states at all, this access is minimal and may not be used to noticeably improve upcoming performance. One reason why access to current performance-related states may be of little use for improving upcoming performance (500 to 1000 ms later) could be down to the difficulty of predicting future internal states from current ones. Although neural correlates of upcoming performance have been identified, these are typically very short-term and their predictive power is very low (see section “Biological underpinnings of variability and performance” below). Although this limited predictive power could be down to technical limitations, we cannot exclude that future performance is to a large extent non-deterministic and therefore largely unpredictable even from within. A conservative interpretation of our results may therefore be that we do have some access to our performance-related internal states, but this access is 1) very limited, 2) rarely spontaneous, and therefore 3) mostly irrelevant to improving future performance.

At first, this interpretation may seem at odds with existing literature on mind wandering, which assumes people can access at least some aspects of their internal fluctuating states. However, our conservative interpretation may link with this literature in a couple of ways. First, limited access would explain why the link between behavioural performance or variability and probe-caught subjective reports of mind wandering is robust but weak. For example, over five different samples, participants who reported being fully mentally ‘zoned out’ from the task only showed an increase of ∼3-7% in variability compared to when they were fully on task (Seli et al., 2013, Laflamme et al., 2018).

Secondly, its lack of spontaneity would match the differences between results from ‘self-caught’ and ‘probe-caught’ methods in the study of mind wandering (see Weinstein, 2017 for a review). Self-caught methods rely on the participant to report each time they are aware they are mind-wandering, whereas probe-caught methods probe participants about their thoughts just prior the probe (which is, as such, always a ‘post-hoc’ judgement), usually at pseudo-random times during the task. The self-caught method is generally not preferred, because participants often do not catch their own deteriorated states of performance (Franklin, Smallwood & Schooler, 2011; Schooler, Reichle & Halpern, 2004). Within the mind wandering literature, this inability to self-catch mind wandering has been explained by a reduction of ‘meta-awareness’ such that if one is mind wandering, and performance is reduced due to a loss of attentional resources, one’s meta-awareness of the mind wandering and deteriorated performance is also reduced. Although mind wandering is a mental states and therefore requires some form of awareness (see Introduction), in these cases, the awareness may be ‘post-hoc’. This inability would then relate to our third point: that our (marginal) access may be no help in improving future performance.

To draw a parallel between our findings and the mind wandering literature, prompting participants would be somewhat similar to the post-error slowing reported in the present study. Similarly, participants are able to access their task-unrelated thoughts when prompted to do so by the experimenter. In contrast, it may be much harder to spontaneously detect mind wandering and other unfavourable states, as would have been required in Experiment 2 in order to use the control when available to prevent errors and very long RT from occurring in the near future.

However, our findings are also theoretically consistent with another (more drastic) interpretation: That we do not have any access to our performance-related inner states. The correlations between behavioural variability and subjective reports of mind wandering could be caused by a third variable that underlies both, but this variable may be fully opaque to us. As often suggested in this literature, such internal states could be related to the activation in the default mode network (Christoff, Gordon, Smallwood, Smith & Schooler, 2009; Mason et al., 2007) or in task-related networks (such as the dorsal attention network; Corbetta, Patel & Shulman, 2008), or the anticorrelation between them (Kelly, Uddin, Biswal, Castellanos & Milham, 2008). As such, behavioural variability or poor performance may not be a direct consequence of mind wandering, but both would likely co-occur in time. Likewise, good performance may co-occur (more often than not) with task-related thoughts or the feeling of being ready, which would also lead to positive associations between subjective reports and behaviour.

The idea that mind wandering may not directly cause poor performance appears at first to contradict previous accounts (eg. “mind wandering influences ongoing primary-task performance”, Laflamme et al. 2018, p.1). However, such accounts may reflect the functional processes that underlie the construct of mind wandering. Previous studies have suggested that mind wandering contains a high proportion of self-oriented thoughts, and seems to play a role in future and autobiographical planning (Baird, Smallwood & Schooler, 2011; D’Argembeau, Renaud & van der Linden, 2011). In this light, mind wandering has been described as a somewhat economical phenomenon: Since the task does not seem to require a high amount of ‘mental/cognitive resources’, they may instead be used to solve self-oriented problems in the meantime. This description is consistent with the findings of Ward & Wegner (2013), who contrasted the construct of mind wandering to that of ‘mind blanking’ (referring to a mental state in which one is void of all thoughts, both task-related and task-unrelated). They concluded participants seemed to find it easier to catch their own mind blanking compared to their mind wandering – and they suggested that because mind wandering may have beneficial components, it is not always necessary to ‘snap out of it’.

To summarise, our findings are consistent with both conservative (people have some access to their internal fluctuating states, but very marginally and typically irrelevant) and extreme interpretations (people have no access to their internal fluctuating states at all). While these interpretations seem at odds with common assumptions, both accounts are reconcilable with current literature.

## Motivation

It could be argued that our results may also be explained by a lack of motivation in our participants. If their motivation was indeed limited to going through the experiment as fast or effortlessly as possible, our participants may not have had the *will* to access performance-relevant information in order to improve their performance. Although we cannot reject that motivation played some role in our results, this interpretation is unlikely to explain our data. In Experiment 1, participants reported high levels of internal motivation (or were otherwise excluded). In Experiment 2, although the task was more boring, our participants were all paid and were mostly postgraduate students who were highly familiar with psychological testing – and as such, expected to show high intrinsic motivation. Moreover, not only their ITIs were twice more variable than would be expected if they simply initiated the next trial as swiftly as possible, but they also massively slowed down following an error (around 40% increase in ITI on average).

We noted that self-paced ITIs showed large individual differences and wondered if these could provide a key to why the control appeared useful to some participants and detrimental to others, resulting in no overall improvement. However, additional between-subject analyses (not presented in paper) did not reveal links between any of the three ITI-characteristics (variability, temporal dependency, and post-error slowing) and the improvement in performance between the Self-paced condition and each of the forced-paced conditions. When instead looking at mean ITI, there was a consistent *negative* relationship with the improvement in performance across all three forced-paced condition: Participants who had a *shorter* mean ITI showed more improvement. As our within-subject analysis showed that longer ITIs may be markers of an overall poor mode of responding – they are followed by poorer rather than better performance, both findings could reflect that good participants indeed show less of these poor modes.

Importantly, our conclusions did not depend on the task being boring or engaging, as both showed similar results. The absence of a benefit of control thus does not seem to rely on motivation.

### Changes in performance versus changes in strategy

To help us resolve ambiguities in the outcome of two of our empirical tests, EZ-diffusion model parameters were calculated for each condition (Wagenmakers et al., 2007). The first ambiguity regarded the interpretation of co-occurring RT increase and % error decrease in two of the forced-paced conditions (Replay and Shuffled Replay) in the action-oriented task. EZ-Diffusion model attributed this difference to higher boundary separation in these conditions, compared to the Self-paced and the Fixed conditions. While a change in drift rate is commonly interpreted as a change in processing efficiency, changes in boundary separation are typically interpreted as strategic changes in caution, i.e. speed-accuracy trade-off, leading us to conclude that there was no improvement in performance in the Self-paced condition. Below we discuss two counterarguments to this interpretation.

First, one could argue that the reduced boundary separations in Self-paced compared to Replay or Shuffled Replay still reflect an active effort of participants to change performance, even if it did not result in a ‘true improvement’. However, if anything, it seems more likely that active effort to change behaviour will lead to increased boundary separation instead, as participants would be more aware of their accuracy than their speed (especially on a milliseconds scale) – i.e., it seems more concrete to aim at ‘zero errors’ than at ‘reducing speed by 50 ms’. In contrast, our results showed that boundary separation was lower in the Self-paced condition. Furthermore, this was very similar to the Fixed condition, and therefore any adjustment is not specific to the Self-paced condition.

Second, we used a simplified version of the drift diffusion model, as it has been shown to be more powerful in detecting effects in drift rate and boundary separation than more complex variants (van Ravenzwaaij, Donkin & Vandekerchove, 2017; van Ravenzwaaij & Oberauer, 2009). Importantly though, this variant does not include a parameter capturing variability in drift rates across trials. Both drift rate and drift rate variability have been associated with self-reported mind wandering, but variability was a stronger predictor (McVay & Kane, 2012, though they used a linear-ballistic accumulator model; LBA). This makes sense when considering that variability is captured by both very long RT and very short RT – the combination of which has a larger effect on variance than on mean performance. A simulation study by van Ravenzwaaij & Oberauer (2009) reported that when data are generated from the LBA and fit with the EZ-diffusion model, increased drift rate variability in the generating (LBA) model is negatively correlated with EZ’s drift rate estimates, and positively correlated with EZ’s estimates of boundary separation. One could question then if the decreased boundary separation in Self-paced actually reflects participants using control to reduce variability in their processing rates. However, data generation with a drift diffusion model shows that such a reduction in drift rate variability would lead to reduced error rates in the Self-paced relative to the Replay and Shuffled Replay conditions, whereas our results show an increase. As such, our results are more consistent with a change in boundary separation.

Altogether, the reported differences are more likely caused by predictability of target onset than by control (see Table 2 for an overview). As such, knowing the time of trial onset seems to lead to a less cautious response pattern (lower decision threshold), in which speed is emphasised over accuracy, with no overall change in processing efficiency, which is in line with previous work (Miller, Sproesser & Ulrich, 2008).

### Changes in non-decision times

The second ambiguity in our results was the smaller number of very short RT and the subsequent decreased variability in the Self-paced perception-oriented task. The EZ-model suggests that, in the perception-oriented task only, non-decision times were higher for self-paced compared to the three control conditions, suggesting slower sensory or motor processes (Wagenmakers et al., 2007). It is unlikely that motor output time would be the cause, as this would be expected to be the same across tasks and conditions. One possibility is that the additional action in the self-paced condition interfered with the sensory processes, but not enough to give hindrance when the stimuli are easy to see.

In any case, unlike in Experiment 2, Experiment 1 did not have an additional action in the self-paced condition. Instead, the Fixed-paced condition in Experiment 1 featured tones, while the Self-paced condition did not. Despite these differences, we found the same results over the experiments; in Experiment 1, the forced-paced condition featured an ‘additional stimulus’, while in Experiment 2, the Self-paced condition featured an ‘additional action’, but results always favoured Hypothesis 2 over Hypothesis 1.

### Routines and practice in sports psychology

After finishing the darts game in Experiment 1, our participants were informally asked if they used any strategies in the Self-paced condition. Many of the participants reported that they threw *“When I felt like it”* or *“When I felt ready for it”*. However, in light of our results, the behavioural relevance of this feeling remains unclear.

While the idea of ‘mentally feeling ready to perform the action’ may seem intuitive to laymen, the area of sports psychology has mainly focused instead on (pre-performance/during-performance) routines as well as the consistency of these routines as means to improve performance (for reviews, see for example: Cohn, 1990; Singer, 2002; Cotterill, 2010). The key element of these routines is training automaticity as a way to enhance task attention and to decrease focus to external distractions.

It should be noted that this ‘automaticity’ may still entail the involvement of cognitive processes, and that sportsmen require mental flexibility for their performance (see Toner, Montero & Moran, 2015 for a review). However, the focus of these cognitive processes does not appear to be on one’s own inner state, but rather with handling relevant environmental states (e.g., the amount and direction of wind when striking with a golf club) or with improving flows in their movement (while aiming to improve one’s skill level). As such, internally-driven variability in sports is often described from a perspective of sensorimotor control; resulting from sources as movement timing and trajectory (e.g., Smeets et al., 2002) rather than from attentional fluctuations. Previous research has found benefits of ‘external foci of attention’ (e.g., focusing on the darts board) over ‘internal foci of attention’ (e.g., focusing on one’s own movement of the arm), both on performance as well as on pre-performance (neuro)physiological states (Marchant, Clough & Crawshaw, 2007; Marchant, Clough, Crawshaw & Levy, 2009; Neumann & Piercy, 2013; Radlo, Steinberg, Singer, Barba & Melnikov, 2002).

In other words, while sports psychology has an interest in reducing variability and creating the most optimal pre-performance state, their interest does not seem to lie in ‘reading inner states’, but rather in training repetitive, automatic, and externally-focused states. Compared to Experiment 1, this type of training would be more similar to the Forced-paced than the Self-paced condition. Interestingly, these ‘repetitive states’ also appears in other aspects of training. Within sports literature, emphasis is put on the consistency of optimal physical movements (for example, consistency in throwing in darts, Brenner, van Dam, Berkhout & Smeets, 2012; Smeets et al., 2002, or in golf, see Langdown, Bridge & Li, 2012 for a review).

One may wonder if the skill level in darts of our participants played a role in Experiment 1, and whether professional darts players *would* be able to use the control effectively. Only one of the twenty-one subjects reported playing darts about twice a week, while all the other subjects had played it a few times a year or less. Therefore, it was not possible to test the effect of skill level in our data (though the scores of the experienced participant did not seem to display a diverging pattern). However, it is important to note that, if anything, the largest numerical differences between the Self-paced and Forced-paced conditions took place in the first block, when participants may still be getting used to the rhythm of the Forced-paced condition. At the later blocks (especially block 4 and 5), the conditions are most similar, suggesting that practice makes the conditions more similar, not less.

### Training access to internal states?

The possible influence of skills and practice on performance leads us to a larger question: To what extent is it possible to train access to our own internal states? One field of research relevant here is mindfulness (meditation) training. Within this literature, people may be trained to be more mindful of their internal states – and as such, may be trained to improve their attention and performance (Brown & Ryan, 2003; Wells, 2005; Zeidan, Johnson, Diamond, David & Goolkasian, 2010) and “tame mind wandering” (Morrison, Goolsarran, Rogers & Jha, 2014; Mrazek, Franklin, Phillips, Baird & Schooler, 2013). However, reported effects tend to be moderate – for instance, Morrison et al., 2014 reported a reduction of ∼8.5% in variability after a seven-hour training over seven weeks. Like attention and mind wandering, mindfulness is a very broad concept (Bergomi, Tschacher & Kupper, 2013) and could refer to a multitude of mechanisms. Furthermore, mindfulness is difficult to capture in an experimental study set-up (for instance, when picking participants or when designing a control condition). Outside the mindfulness/meditation literature, Baldwin et al. (2017) found an increase in participants’ own awareness of mind wandering over the course of a five-day experiment. However, due to the highly repetitive nature of the task, it is plausible that participants just allowed themselves to deliberately mind wander more throughout the sessions.

### Temporal dependency

In Experiment 2, we found strong evidence for temporal dependency in both RT and ITI. All comparisons between the self-paced condition and the forced-paced conditions showed evidence against a reduction in temporal dependency, with one exception: The slopes of the Fixed condition in the action-oriented task. Although the evidence was weak and did not replicate in the perception-oriented task, this finding is similar to that of Kelly et al. (2001). Interestingly, they mention that *“self-pacing means that the system is sampled at irregular intervals in real time, violating the assumptions of most dynamical analyses”* (p.824), and propose that conditions with fixed pacing could be more suited for measuring temporal dependency. While their experiment only used forced-paced conditions with a fixed interval, we also used conditions that are forced-paced yet variable. On these variable conditions, we found no differences compared to the Self-paced condition. This is consistent with the notion that conditions with variable intervals may interfere with the measurement of temporal dependency, although this happens irrespective of whether these are self-paced or not.

The temporal dependency in the RT and ITI may suggest that these are coupled to underlying fluctuating states. One commonly mentioned state is ‘attention’, also thought to fluctuate over time and to influence performance. However, Wagenmakers et al. (2004) noted that it is unclear *how* attention would cause the specific temporal patterns common in empirical data. Alternatively, temporal dependencies could be caused by the combination of a number of different processes with varying timescales. This is in line with findings that variability is underpinned by a number of biological processes, all with varying time scales (see below section on *Biological underpinnings of variability and performance*).

### Biological underpinnings of variability and performance

Above, we referred to internal states that may underlie both behavioural performance and variability. These may be reflected in fluctuations in the DMN, task-related networks, and the episodic memory network, which have been often associated with mind wandering (for a meta-analysis, see Fox, Spreng, Ellamil, Andrews-Hanna & Christoff, 2015; for reviews, see Christoff, 2012; Smallwood, Brown, Baird & Schooler, 2012). Indeed, slow rhythms (∼.05 Hz) in BOLD activity within the DMN have been associated with reaction time variability (Weissman, Roberts, Visscher & Woldorff, 2006). Variability in performance on detection or discrimination tasks has been related to oscillatory activity using EEG or MEG, in particular alpha (8-12Hz) power and phase, but also to beta and gamma power (Busch, Dubois & VanRullen, 2009; Van Dijk, Schoffelen, Oostenveld & Jensen, 2008; Drewes & VanRullen, 2011; Ergenoglu et al., 2004; Hanslmayr et al., 2007; Rihs, Michel & Thut, 2007; Romei et al., 2008; Romei et al., 2010; De Graaf et al., 2015; Thut, Nietzel, Brandt & Pascual-Leone, 2006; VanRullen, Busch, Drewes & Dubois, 2011; Bompas et al. 2015). Interestingly, a recent study has shown that spontaneous fluctuations in alpha rhythms are partially locked to slow rhythms (∼.05 Hz) in the stomach, with so called ‘gastric phase’ explaining about 8% of the variance in alpha (Richter, Babo-Rebelo, Schwartz & Tallon-Baudry, 2017). Heartbeat has also been found to play a role in variability in accuracy, such that detection performance is worse if stimuli are presented synchronous with one’s heart beat (Salomon et al., 2016). It is largely unknown to what extent spontaneous variability within these sources could be accessible to consciousness. Whether this knowledge could be used to improve behaviour also heavily relies on the time scale at which this variability unfolds.

### Variability – a beneficial characteristic?

Within many contexts, both in our daily lives as well as in the laboratory, it may be tempting to see variability as a hindering by-product of a lack of attention which we would like to reduce as much as possible. However, variability may not necessarily be negative. Indeed, variability may ensure our behaviour is not entirely predictable to our preys and predators (Carpenter, 1999), and may facilitate exploration and novel behaviour (Shahan & Chase, 2002; see Sternad, 2018 for a review). Furthermore, variability and the resulting unpredictability of our behaviours are key to discussions about our sense of agency and beliefs of free will (see for examples: Brembs, 2011; Haggard, 2008; Koch, 2009; Tse, 2013). This is reflected in models of decision, where noise plays a crucial role (Bogacz, Brown, Moehlis, Holmes & Cohen, 2006; Bompas, Hedge & Sumner, 2017; Bompas & Sumner, 2011).

Importantly, variability is not limited to behaviour, but present throughout all levels of our central nervous system, even in very short-term fluctuations such as the firing of action potentials within a single neuron (with random noise contributing to whether the action potential will be initiated) and subsequent variability in post-synaptic response (which may similarly be affected by ‘synaptic background noise’). Such fluctuations throughout various levels in our nervous system may affect trial-to-trial variability. This variability does not only occur as a response to external stimuli, but also in the absence thereof, as an intrinsic characteristic of the system. It has been argued that randomness is important for the functioning of the nervous system, rather than something that needs to be reduced or ‘overcome’ (Ermentrout, Galán & Urban, 2008; Faisal, Selen & Wolpert, 2008). All in all, variability appears to be an intrinsic and fundamental property, and as such, a large proportion of it may be not reducible at all.

## Conclusion

Intuitively, it seems reasonable to think that people have some access to their own fluctuating performance-relevant inner states, and that they can use this information to improve their performance. In two separate experiments and across a series of empirical tests, we found repeated evidence against most predictions derived from this intuition. We found that, even though people varied the time they initiated a trial and reported that they threw darts only when ready, they were unable to improve their performance or reduce their variability, even when highly motivated to do so. Altogether, this suggests that if people have any access to their own inner states at all, this access is limited and not a key determinant for upcoming performance.

### Context paragraph

This research came about through an interest in the reducibility of intra-individual variability – a question that may be approached from two distinct lines of research. From one perspective, variability is perceived as a negative consequence of attentional lapses, which should be reduced as much as possible. Empirical evidence comes from studies on mind wandering, showing that reduced attention correlates with higher variability, and on attention-focused training (e.g., mindfulness) that lead to reduced variability. However, reported effects are often weak and may be sensitive to publication biases. Still, the perspective that we can improve our performance by voluntarily focusing on the task seems highly intuitive. From the second perspective though, variability results from an intrinsic and necessary property of our nervous system, arising from a multitude of inaccessible sources and may therefore not be reducible voluntarily. To deliver the first empirical test contrasting these two perspectives, we investigated whether participants could improve performance when offered some control over the experiment. Importantly, we were not theoretically biased and were determined to report either outcome. To provide a comprehensive answer, we used multiple paradigms and statistical tests – which all failed to support the intuitive framework.

## Acknowledgments

This work was partly funded by the ESRC (ES/K002325/1). Funding sources had no involvement at any stage of the reported research. We are grateful to Simon Farrell for sharing his code for the spectral analysis. We would like to thank Nia Paddison Rees and Joyce Ira Go, who helped collecting the data for Experiment 1, and Natasha Harris, Humairaa Uddin, and Sarah Lethbridge, who helped collecting the data for Experiment 2.

## Supplementary Materials

In order to compare temporal dependencies across conditions, we first tested whether our RT and ITI measures actually contained temporal dependencies. Bayesian One Sample one-sided t-tests were used to test if the participants’ autocorrelations at lag one (AC1) and the slopes of the linearly fitted power spectra were statistically higher than zero (see Supplementary Table 1 for the corresponding BF). Out of ten data series (RT on four conditions plus self-paced ITIs, for two tasks), eight showed evidence for temporal dependencies while two did not (Self-paced and Shuffled replay RTs in the action-oriented task). Evidence for dependencies were consistent across both measures.

**Supplementary Table 1.** Bayes’ Factors for the presence of temporal dependencies in the RT and self-paced ITIs, tested against two different measures: a positive autocorrelation function at lag 1 (AC1), and slope of the power spectrum.

| **Test** | **Action-oriented** |  | **Perception-oriented** |  |
| --- | --- | --- | --- | --- |
|  | **AC1** | **Slope** | **AC1** | **slope** |
| SP<sub>RT</sub> | 1.15 | .37 | 2499.58 | 46.30 |
| F<sub>RT</sub> | 6.53 | 18.26 | 601.65 | 152.45 |
| R<sub>RT</sub> | 11.31 | 1.06 | 359.38 | 416.93 |
| SR<sub>RT</sub> | .50 | .23 | 7.15 | 3.17 |
| SP<sub>ITI</sub> | 22.06 | 5.433e+6 | 24.54 | 6351.91 |

## References

Baird, B., Smallwood, J., & Schooler, J. W. (2011). Back to the future: Autobiographical planning and the functionality of mind-wandering. Consciousness and Cognition, 20(4), 1604–1611. https://doi.org/10.1016/j.concog.2011.08.007

Baldwin, C. L., Roberts, D. M., Barragan, D., Lee, J. D., Lerner, N., & Higgins, J. S. (2017). Detecting and Quantifying Mind Wandering during Simulated Driving. Frontiers in Human Neuroscience, 11. https://doi.org/10.3389/fnhum.2017.00406

Bergomi, C., Tschacher, W., & Kupper, Z. (2013). The Assessment of Mindfulness with Self-Report Measures: Existing Scales and Open Issues. Mindfulness, 4(3), 191–202. https://doi.org/10.1007/s12671-012-0110-9

Bogacz, R., Brown, E., Moehlis, J., Holmes, P., & Cohen, J. D. (2006). The physics of optimal decision making: A formal analysis of models of performance in two-alternative forced-choice tasks. Psychological Review, 113(4), 700–765. https://doi.org/10.1037/0033-295X.113.4.700

Bompas, A., & Sumner, P. (2011). Saccadic Inhibition Reveals the Timing of Automatic and Voluntary Signals in the Human Brain. Journal of Neuroscience, 31(35), 12501–12512. https://doi.org/10.1523/JNEUROSCI.2234-11.2011

Bompas, A., Hedge, C., & Sumner, P. (2017). Speeded saccadic and manual visuo-motor decisions: Distinct processes but same principles. Cognitive Psychology, 94, 26–52. https://doi.org/10.1016/j.cogpsych.2017.02.002

Bompas, A., Sumner, P., Muthumumaraswamy, S. D., Singh, K. D., & Gilchrist, I. D. (2015). The contribution of pre-stimulus neural oscillatory activity to spontaneous response time variability. NeuroImage, 107, 34–45. https://doi.org/10.1016/j.neuroimage.2014.11.057

Brainard, D. H. (1997). The Psychophysics Toolbox. Spatial Vision, 10(4), 433–436. https://doi.org/10.1163/156856897X00357

Brembs, B. (2011). Towards a scientific concept of free will as a biological trait: spontaneous actions and decision-making in invertebrates. Proceedings of the Royal Society B: Biological Sciences, 278(1707), 930–939. https://doi.org/10.1098/rspb.2010.2325

Brenner, E., van Dam, M., Berkhout, S., & Smeets, J. B. J. (2012). Timing the moment of impact in fast human movements. Acta Psychologica, 141, 104–111. https://doi.org/10.1016/j.actpsy.2012.07.002

Brown, K. W., & Ryan, R. M. (2003). The benefits of being present: Mindfulness and its role in psychological well-being. Journal of Personality and Social Psychology, 84(4), 822–848. https://doi.org/10.1037/0022-3514.84.4.822

Busch, N. A., Dubois, J., & VanRullen, R. (2009). The Phase of Ongoing EEG Oscillations Predicts Visual Perception. Journal of Neuroscience, 29(24), 7869–7876. https://doi.org/10.1523/JNEUROSCI.0113-09.2009

Carpenter, R. H. S. (1999). A neural mechanism that randomises behaviour. Journal of Consciousness Studies, 6(1), 13–22.

Cheyne, A. J., Solman, G. J. F., Carriere, J. S. A., & Smilek, D. (2009). Anatomy of an error: A bidirectional state model of task engagement/disengagement and attention-related errors. Cognition, 111(1), 98–113. https://doi.org/10.1016/j.cognition.2008.12.009

Cheyne, J. A., Carriere, J. S. A., & Smilek, D. (2006). Absent-mindedness: Lapses of conscious awareness and everyday cognitive failures. Consciousness and Cognition, 15(3), 578–592. https://doi.org/10.1016/j.concog.2005.11.009

Christoff, K. (2012). Undirected thought: Neural determinants and correlates. Brain Research, 1428, 51–59. https://doi.org/10.1016/j.brainres.2011.09.060

Christoff, K., Gordon, A. M., Smallwood, J., Smith, R., & Schooler, J. W. (2009). Experience sampling during fMRI reveals default network and executive system contributions to mind wandering. Proceedings of the National Academy of Sciences, 106(21), 8719–8724. https://doi.org/10.1073pnas.0900234106

Cohen, M. R., & Maunsell, J. H. R. (2009). Attention improves performance primarily by reducing interneuronal correlations. Nature Neuroscience, 12(12), 1594–1600. https://doi.org/10.1038/nn.2439

Cohn, P. J. (1990). Preperformance Routines in Sport: Theoretical Support and Practical Applications. The Sport Psychologist, 4(3), 301–312. https://doi.org/10.1123/tsp.4.3.301

Corbetta, M., Patel, G., & Shulman, G. L. (2008). The Reorienting System of the Human Brain: From Environment to Theory of Mind. Neuron, 58(3), 306–324. https://doi.org/10.1016/j.neuron.2008.04.017

Cotterill, S. (2010). Pre-performance routines in sport: current understanding and future directions. International Review of Sport and Exercise Psychology, 3(2), 132–153. https://doi.org/10.1080/1750984X.2010.488269

Cubitt, R. P., Starmer, C., & Sugden, R. (1998). On the validity of the random lottery incentive system. Experimental Economics, 1(2), 115–131. https://doi.org/10.1007/BF01669298

Danielmeier, C., & Ullsperger, M. (2011). Post-Error Adjustments. Frontiers in Psychology, 2, 233. https://doi.org/10.3389/fpsyg.2011.00233

D’Argembeau, A., Renaud, O., & Linden, M. V. der. (2011). Frequency, characteristics and functions of future-oriented thoughts in daily life. Applied Cognitive Psychology, 25(1), 96–103. https://doi.org/10.1002/acp.1647

De Graaf, T. A. de, Gross, J., Paterson, G., Rusch, T., Sack, A. T., & Thut, G. (2013). Alpha-Band Rhythms in Visual Task Performance: Phase-Locking by Rhythmic Sensory Stimulation. PLOS ONE, 8(3), e60035. https://doi.org/10.1371/journal.pone.0060035

Drewes, J., & VanRullen, R. (2011). This Is the Rhythm of Your Eyes: The Phase of Ongoing Electroencephalogram Oscillations Modulates Saccadic Reaction Time. Journal of Neuroscience, 31(12), 4698–4708. https://doi.org/10.1523/JNEUROSCI.4795-10.2011

Dutilh, G., van Ravenzwaaij, D., Nieuwenhuis, S., van der Maas, H. L. J., Forstmann, U., & Wagenmakers, E.-J. (2012). How to measure post-error slowing: A confound and a simple solution. Journal of Mathematical Psychology, 56(3), 208–216. https://doi.org/10.1016/j.jmp.2012.04.001

Dutilh, G., Vandekerckhove, J., Forstmann, B. U., Keuleers, E., Brysbaert, M., & Wagenmakers, E.-J. (2012). Testing theories of post-error slowing. Attention, Perception, & Psychophysics, 74(2), 454–465. https://doi.org/10.3758/s13414-011-0243-2

Ergenoglu, T., Demiralp, T., Bayraktaroglu, Z., Ergen, M., Beydagi, H., & Uresin, Y. (2004). Alpha rhythm of the EEG modulates visual detection performance in humans. Cognitive Brain Research, 20(3), 376–383. https://doi.org/10.1016/j.cogbrainres.2004.03.009

Ermentrout, G. B., Galán, R. F., & Urban, N. N. (2008). Reliability, synchrony and noise. Trends in Neurosciences, 31(8), 428–434. https://doi.org/10.1016/j.tins.2008.06.002

Farrell, S., Wagenmakers, E.-J., & Ratcliff, R. (2006). 1/f noise in human cognition: Is it ubiquitous, and what does it mean? Psychonomic bulletin & review, 13(4), 737–741. https://doi.org/10.3758/BF03193989

Faisal, A. A., Selen, L. P. J., & Wolpert, D. M. (2008). Noise in the nervous system. Nature Reviews Neuroscience, 9(4), 292–303. https://doi.org/10.1038/nrn2258

Fox, K. C. R., Spreng, R. N., Ellamil, M., Andrews-Hanna, J. R., & Christoff, K. (2015). The wandering brain: Meta-analysis of functional neuroimaging studies of mind-wandering and related spontaneous thought processes. NeuroImage, 111, 611–621. https://doi.org/10.1016/j.neuroimage.2015.02.039

Franklin, M. S., Smallwood, J., & Schooler, J. W. (2011). Catching the mind in flight: Using behavioral indices to detect mindless reading in real time. Psychonomic Bulletin & Review, 18(5), 992–997. https://doi.org/10.3758/s13423-011-0109-6

Giambra, L. M. (1995). A Laboratory Method for Investigating Influences on Switching Attention to Task-Unrelated Imagery and Thought. Consciousness and Cognition, 4(1), 1–21. https://doi.org/10.1006/ccog.1995.1001

Gilden, D. L. (2001). Cognitive Emissions of 1/f Noise. Psychological Review, 108(1), 33–56. https://doi.org/10.I037//0033-295X.108.1.33

Gilden, D. L., & Wilson, S. G. (1995). Streaks in skilled performance. Psychonomic Bulletin & Review, 2(2), 260–265. https://doi.org/10.3758/BF03210967

Haggard, P. (2008). Human volition: towards a neuroscience of will. Nature Reviews Neuroscience, 9(12), 934–946. https://doi.org/10.1038/nrn2497

Hanslmayr, S., Aslan, A., Staudigl, T., Klimesch, W., Herrmann, C. S., & Bäuml, K.-H. (2007). Prestimulus oscillations predict visual perception performance between and within subjects. NeuroImage, 37(4), 1465–1473. https://doi.org/10.1016/j.neuroimage.2007.07.011

Huber, M. E., Kuznetsov, N., & Sternad, D. (2016). Persistence of reduced neuromotor noise in long-term motor skill learning. Journal of Neurophysiology, 116(6), 2922–2935. https://doi.org/10.1152/jn.00263.2016

JASP Team (2017). JASP (Version 0.8.5).

Kelly, A., Heathcote, A., Heath, R., & Longstaff, M. (2001). Response-Time Dynamics: Evidence for Linear and Low-Dimensional Nonlinear Structure in Human Choice Sequences. The Quarterly Journal of Experimental Psychology Section A, 54(3), 805–840. https://doi.org/10.1080/713755987

Kelly, A. M. C., Uddin, L. Q., Biswal, B. B., Castellanos, F. X., & Milham, M. P. (2008). Competition between functional brain networks mediates behavioral variability. NeuroImage, 39(1), 527–537. https://doi.org/10.1016/j.neuroimage.2007.08.008

Kleiner, M., Brainard, D. H., & Pelli, D. (2007). What’s new in Psychtoolbox-3? Perception, 36(14), 1–16. https://doi.org/10.1068/v070821

Koch, C. (2009). Free Will, Physics, Biology, and the Brain. In N. Murphy, G. R. R. Ellis, & T. O’Connor (Red.), Downward Causation and the Neurobiology of Free Will (pp. 31–52). Springer, Berlin, Heidelberg. https://doi.org/10.1007/978-3-642-03205-9_2

Kofler, M. J., Rapport, M. D., Sarver, D. E., Raiker, J. S., Orban, S. A., Friedman, L. M., & Kolomeyer, E. G. (2013). Reaction time variability in ADHD: a metaanalytic review of 319 studies. Clinical Psychology Review, 33(6), 795–811. https://doi.org/10.1016/j.cpr.2013.06.001

Kontsevich, L. L., & Tyler, C. W. (1999). Bayesian adaptive estimation of psychometric slope and threshold. Vision Research, 39(16), 2729–2737. https://doi.org/10.1016/S0042-6989(98)00285-5

Langdown, B. L., Bridge, M. & Li, F.-X. (2012). Movement variability in the golf swing. Sports Biomechanics, 11(2), 273–287. https://doi.org/10.1080/14763141.2011.650187

Laflamme, P., Seli, P., & Smilek, D. (2018). Validating a visual version of the metronome response task. Behavior Research Methods, 1–12. https://doi.org/10.3758/s13428-018-1020-0

Ledberg, A., Montagnini, A., Coppola, R., & Bressler, S. L. (2012). Reduced Variability of Ongoing and Evoked Cortical Activity Leads to Improved Behavioral Performance. PLOS ONE, 7(8), e43166. https://doi.org/10.1371/journal.pone.0043166

Marchant, D. C., Clough, P. J., & Crawshaw, M. (2007). The effects of attentional focusing strategies on novice dart throwing performance and Their task experiences. International Journal of Sport and Exercise Psychology, 5(3), 291–303. https://doi.org/10.1080/1612197X.2007.9671837

Marchant, D. C., Clough, P. J., Crawshaw, M., & Levy, A. (2009). Novice motor skill performance and task experience is influenced by attentional focusing instructions and instruction preferences. International Journal of Sport and Exercise Psychology, 7(4), 488–502. https://doi.org/10.1080/1612197X.2009.9671921

Marom, S., & Wallach, A. (2011). Relational dynamics in perception: impacts on trial-to-trial variation. Frontiers in Computational Neuroscience, 5, 16. https://doi.org/10.3389/fncom.2011.00016

Mason, M. F., Norton, M. I., Van Horn, J. D., Wegner, D. M., Grafton, S. T., & Macrae, N. (2007). Wandering Minds: The Default Network and Stimulus-Independent Thought. Science, 315(5810), 393–395. https://doi.org/10.1126/science.1131295

Masquelier, T. (2013). Neural variability, or lack thereof. Frontiers in Computational Neuroscience, 7, 1–7. https://doi.org/10.3389/fncom.2013.00007

McAuley, E., Wraith, S., & Duncan, T. E. (1991). Self-Efficacy, Perceptions of Success, and Intrinsic Motivation for Exercise1. Journal of Applied Social Psychology, 21(2), 139–155. https://doi.org/10.1111/j.1559-1816.1991.tb00493.x

McVay, J. C., & Kane, M. J. (2012). Drifting from Slow to “D’oh!” Working Memory Capacity and Mind Wandering Predict Extreme Reaction Times and Executive-Control Errors. Journal of Experimental Psychology. Learning, Memory, and Cognition, 38(3), 525–549. https://doi.org/10.1037/a0025896

Miller, J., Sproesser, G., & Ulrich, R. (2008). Constant versus variable response signal delays in speed-accuracy trade-offs: Effects of advance preparation for processing time. Perception & Psychophysics, 70(5), 878–886. https://doi.org/10.3758/PP.70.5.878

Mitchell, J. F., Sundberg, K. A., & Reynolds, J. H. (2007). Differential Attention-Dependent Response Modulation across Cell Classes in Macaque Visual Area V4. Neuron, 55(1), 131–141. https://doi.org/10.1016/j.neuron.2007.06.018

Morrison, A. B., Goolsarran, M., Rogers, S. L., & Jha, A. P. (2014). Taming a wandering attention: short-form mindfulness training in student cohorts. Frontiers in Human Neuroscience, 7. https://doi.org/10.3389/fnhum.2013.00897

Mrazek, M. D., Smallwood, J., & Schooler, J. W. (2012). Mindfulness and mind-wandering: finding convergence through opposing constructs. Emotion, 12(3), 442–448. https://doi.org/10.1037/a0026678

Neumann, D. L., & Piercy, A. (2013). The Effect of Different Attentional Strategies on Physiological and Psychological States During Running. Australian Psychologist, 48(5), 329–337. https://doi.org/10.1111/ap.12015

Pelli, D. G. (1997). The VideoToolbox software for visual psychophysics: Transforming numbers into movies. Spatial Vision, 10(4), 437–442. https://doi.org/10.1163/156856897X00366

R Core Team (2013). R: A language and environment for statistical computing. R Foundation or Statistical Computing, Vienna, Austria. URL http://www.R-project.org

Rabbitt, P. M. (1966). Errors and error correction in choice-response tasks. Journal of Experimental Psychology, 71(2), 264–272. https://doi.org/10.1037/h0022853

Radlo, S. J., Steinberg, G. M., Singer, R. N., Barba, D. A., & Melnikov, A. (2002). The influence of an attentional focus strategy on alpha brain wave activity, heart rate, and dart-throwing performance. International Journal of Sport Psychology, 33(2), 205–217.

Ratcliff, R. (1978). A theory of memory retrieval. Psychological Review, 85(2), 59–108. https://doi.org/10.1037/0033-295X.85.2.59

Richter, C. G., Babo-Rebelo, M., Schwartz, D., & Tallon-Baudry, C. (2017). Phase-amplitude coupling at the organism level: The amplitude of spontaneous alpha rhythm fluctuations varies with the phase of the infra-slow gastric basal rhythm. NeuroImage, 146, 951–958. https://doi.org/10.1016/j.neuroimage.2016.08.043

Rihs, T. A., Michel, C. M., & Thut, G. (2007). Mechanisms of selective inhibition in visual spatial attention are indexed by α-band EEG synchronization. European Journal of Neuroscience, 25(2), 603–610. https://doi.org/10.1111/j.1460-9568.2007.05278.x

Romei, V., Gross, J., & Thut, G. (2010). On the Role of Prestimulus Alpha Rhythms over Occipito-Parietal Areas in Visual Input Regulation: Correlation or Causation? Journal of Neuroscience, 30(25), 8692–8697. https://doi.org/10.1523/JNEUROSCI.0160-10.2010

Romei, V., Rihs, T., Brodbeck, V., & Thut, G. (2008). Resting electroencephalogram alpha-power over posterior sites indexes baseline visual cortex excitability: NeuroReport, 19(2), 203–208. https://doi.org/10.1097/WNR.0b013e3282f454c4

Salomon, R., Ronchi, R., Dönz, J., Bello-Ruiz, J., Herbelin, B., Martet, R., Faivre, N., Schaller, K., & Blanke, O. (2016). The Insula Mediates Access to Awareness of Visual Stimuli Presented Synchronously to the Heartbeat. Journal of Neuroscience, 36(18), 5115–5127. https://doi.org/10.1523/JNEUROSCI.4262-15.2016

Schooler, J. W., Reichle, E. D., & Halpern, D. V. (2004). Zoning Out while Reading: Evidence for Dissociations between Experience and Metaconsciousness. In D. T. Levin (Red.), Thinking and seeing: visual metacognition in adults and children. Cambridge, Mass: MIT Press.

Seli, P., Cheyne, J. A., & Smilek, D. (2013). Wandering minds and wavering rhythms: linking mind wandering and behavioral variability. Journal of Experimental Psychology: Human Perception and Performance, 39(1), 1–5. https://doi.org/10.1037/a0030954

Seli, P., Smallwood, J., Cheyne, J. A., & Smilek, D. (2015). On the relation of mind wandering and ADHD symptomatology. Psychonomic Bulletin & Review, 22(3), 629–636. https://doi.org/10.3758/s13423-014-0793-0

Shahan, T. A., & Chase, P. N. (2002). Novelty, stimulus control, and operant variability. The Behavior Analyst, 25(2), 175–190. https://doi.org/10.1007/BF03392056

Shaw, G. A., & Giambra, L. (1993). Task-unrelated thoughts of college students diagnosed as hyperactive in childhood. Developmental Neuropsychology, 9(1), 17–30. https://doi.org/10.1080/87565649309540541

Singer, R. N. (2002). Preperformance State, Routines, and Automaticity: What Does It Take to Realize Expertise in Self-Paced Events? Journal of Sport and Exercise Psychology, 24(4), 359–375. https://doi.org/10.1123/jsep.24.4.359

Smallwood, J., Brown, K., Baird, B., & Schooler, J. W. (2012). Cooperation between the default mode network and the frontal–parietal network in the production of an internal train of thought. Brain Research, 1428, 60–70. https://doi.org/10.1016/j.brainres.2011.03.072

Smeets, J. B. J., Frens, M. A., & Brenner, E. (2002). Throwing darts: timing is not the limiting factor. Experimental Brain Research, 144, 268–274.

Smith, G. (2003). Horseshoe pitchers’ hot hands. Psychonomic Bulletin & Review, 10(3), 753–758. https://doi.org/10.3758/BF03196542

Sternad, D. (2018). It’s not (only) the mean that matters: variability, noise and exploration in skill learning. Current Opinion in Behavioral Sciences, 20, 183–195. https://doi.org/10.1016/j.cobeha.2018.01.004

Stins, J. F., Yaari, G., Wijmer, K., Burger, J. F., & Beek, P. J. (2018). Evidence for Sequential Performance Effects in Professional Darts. Frontiers in Psychology, 9, 591. https://doi.org/10.3389/fpsyg.2018.00591

Tamm, L., Narad, M. E., Antonini, T. N., O’Brien, K. M., Hawk, L. W., & Epstein, J. N. (2012). Reaction Time Variability in ADHD: A Review. Neurotherapeutics, 9(3), 500–508. https://doi.org/10.1007/s13311-012-0138-5

Tauer, J. M., & Harackiewicz, J. M. (2004). The Effects of Cooperation and Competition on Intrinsic Motivation and Performance. Journal of Personality and Social Psychology, 86(6), 849–861. https://doi.org/10.1037/0022-3514.86.6.849

The MathWorks, Inc. (Release 2016a). MATLAB 9. Natick, Massachusetts, United States.

Thomson, D. R., Seli, P., Besner, D., & Smilek, D. (2014). On the link between mind wandering and task performance over time. Consciousness and Cognition, 27, 14–26. https://doi.org/10.1016/j.concog.2014.04.001=

Thut, G., Nietzel, A., Brandt, S. A., & Pascual-Leone, A. (2006). α-Band Electroencephalographic Activity over Occipital Cortex Indexes Visuospatial Attention Bias and Predicts Visual Target Detection. Journal of Neuroscience, 26(37), 9494–9502. https://doi.org/10.1523/JNEUROSCI.0875-06.2006

Toner, J., Montero, B. G., & Moran, A. (2015). Considering the role of cognitive control in expert performance. Phenomenology and the Cognitive Sciences, 14(4), 1127–1144. http://dx.doi.org/10.1007/s11097-014-9407-6

Tse, P. (2013). The Neural Basis of Free Will: Criterial Causation. MIT Press.

Van Beers, R. J., van der Meer, Y., & Veerman, R. M. (2013). What Autocorrelation Tells Us about Motor Variability: Insights from Dart Throwing. Plos ONE 8(5): e64332. doi:10.1371/journal.pone.0064332

Van Dijk, H., Schoffelen, J.-M., Oostenveld, R., & Jensen, O. (2008). Prestimulus oscillatory activity in the alpha band predicts visual discrimination ability. The Journal of Neuroscience: The Official Journal of the Society for Neuroscience, 28(8), 1816–1823. https://doi.org/10.1523/JNEUROSCI.1853-07.2008

van Ravenzwaaij, D., Donkin, C., & Vandekerckhove, J. (2017). The EZ diffusion model provides a powerful test of simple empirical effects. Psychonomic Bulletin & Review, 24(2), 547–556. https://doi.org/10.3758/s13423-016-1081-y

van Ravenzwaaij, D., & Oberauer, K. (2009). How to use the diffusion model: Parameter recovery of three methods: EZ, fast-dm, and DMAT. Journal of Mathematical Psychology, 53(6), 463–473. https://doi.org/10.1016/j.jmp.2009.09.004

VanRullen, R., Busch, N., Drewes, J., & Dubois, J. (2011). Ongoing EEG Phase as a Trial-by-Trial Predictor of Perceptual and Attentional Variability. Frontiers in Psychology, 2, 1–9. https://doi.org/10.3389/fpsyg.2011.00060

Wagenmakers, E.-J., Farrell, S., & Ratcliff, R. (2004). Estimation and interpretation of 1/fα noise in human cognition. Psychonomic Bulletin & Review, 11(4), 579–615. https://doi.org/10.3758/BF03196615

Wagenmakers, E.-J., Van Der Maas, H. L. J., & Grasman, R. P. P. P. (2007). An EZ-diffusion model for response time and accuracy. Psychonomic Bulletin & Review, 14(1), 3–22. https://doi.org/10.3758/BF03194023

Ward, A. F., & Wegner, D. M. (2013). Mind-blanking: When the mind goes away. Frontiers in Psychology, 4. https://doi.org/10.3389/fpsyg.2013.00650

Weinstein, Y. (2017). Mind-wandering, how do I measure thee with probes? Let me count the ways. Behavior Research Methods, 50(2), 642–661. https://doi.org/10.3758/s13428-017-0891-9

Weissman, D. H., Roberts, K. C., Visscher, K. M., & Woldorff, M. G. (2006). The neural bases of momentary lapses in attention. Nature Neuroscience, 9(7), 971–978. https://doi.org/10.1038/nn1727

Wells, A. (2005). Detached Mindfulness In Cognitive Therapy: A Metacognitive Analysis And Ten Techniques. Journal of Rational-Emotive & Cognitive-Behavior Therapy, 23(4), 337–355. https://doi.org/10.1007/s10942-005-0018-6

Wessel, J. R., & Aron, A. R. (2017). On the Globality of Motor Suppression: Unexpected Events and Their Influence on Behavior and Cognition. Neuron, 93(2), 259–280. https://doi.org/10.1016/j.neuron.2016.12.013

Zeidan, F., Johnson, S. K., Diamond, B. J., David, Z., & Goolkasian, P. (2010). Mindfulness meditation improves cognition: Evidence of brief mental training. Consciousness and Cognition, 19(2), 597–605. https://doi.org/10.1016/j.concog.2010.03.014

